# Cortical layer 6b mediates state-dependent changes in brain activity and effects of orexin on waking and sleep

**DOI:** 10.1101/2024.10.26.620399

**Authors:** Elise J. Meijer, Marissa Mueller, Lukas B. Krone, Tomoko Yamagata, Anna Hoerder-Suabedissen, Sian Wilcox, Hannah Alfonsa, Atreyi Chakrabarty, Luiz Guidi, Peter L. Oliver, Vladyslav V. Vyazovskiy, Zoltán Molnár

## Abstract

One of the most distinctive features of the mammalian cerebral cortex is its laminar structure. Of all cortical layers, layer 6b (L6b) is by far the least-studied, despite exhibiting direct sensitivity to orexin and having widespread connectivity, suggesting an important role in regulating cortical oscillations and brain state. We performed chronic electroencephalogram (EEG) recordings in mice in which a subset of L6b neurons was conditionally “silenced”, during undisturbed conditions, after sleep deprivation (SD), and after intracerebroventricular (ICV) administration of orexin. While the total amount of waking and sleep or the response to SD were not altered, L6b-silenced mice showed a slowing of theta frequency (6-9 Hz) during wake and REM sleep, and a marked reduction of total EEG power, especially in NREM sleep. The infusion of orexin A increased wakefulness in both genotypes, but subsequent levels of EEG slow-wave activity during NREM sleep were lower in L6b-silenced animals in the occipital derivation. In summary, these results demonstrate a role for cortical L6b in state-dependent brain oscillations and in the response to orexinergic neurotransmission. Our findings provide new insights in functions of L6b neurons and could inform the understanding of abnormal regulation of brain states in neurodevelopmental and psychiatric disorders.

## Introduction

Layer 6b (L6b) is the earliest generated and deepest layer of the mouse neocortex and is partially derived from the subplate – a critical structure during cortical development which contributes to the formation of intracortical and thalamocortical networks (McConnell et al., 1989; Allendoerfer and Shatz, 1994; Molnár and Blakemore, 1995; Molnár et al., 1998; Feldmeyer, 2023). While evolutionary conservation of L6b neurons and their primate homologue interstitial white matter neurons (IWN) across many species suggests that these cells are not merely remnants of a developmental cell population, their functional significance in the adult brain remains unknown (Kostovic and Rakic, 1980; Reep, 2000; Kostović and Judaš, 2010; Duque et al., 2016; Molnar, 2018; Swiegers et al., 2019, 2021; Bhagwandin et al., 2020).

L6b exhibits widespread anatomical connectivity, implying an integrative role across a range of functional brain networks. L6b receives long-range intracortical inputs, enabling direct communication between distant cortical areas (Hoerder-Suabedissen et al., 2018; Zolnik et al., 2020), while being reciprocally innervated by higher order thalamic nuclei and cortical layer 5 (L5) (Zolnik et al., 2024). Consistent with this notion, it was shown that optogenetic activation of L6b neurons leads to activation of the posterior medial nucleus of the thalamus (Ansorge et al., 2020) and the neocortex (Zolnik et al., 2024). This connectivity is of potential importance for brain state regulation as L5 is the major output layer of the cortex, and is engaged in bidirectional communication with higher order thalamic nuclei (Hoerder-Suabedissen et al., 2018). Recently, several subtypes of L5 pyramidal neurons have been shown to play important roles in distinct aspects of sleep-wake regulation (Krone et al., 2021, 2025; Hong et al., 2023; Honjoh et al., 2025; Wasilczuk et al., 2025). Together with the established role of L5 pyramidal neurons in the regulation of oscillatory dynamics with functional relevance for sleep (Sanchez-Vives and McCormick, 2000; Seibt et al., 2017) and anaesthesia (Bharioke et al., 2022), the projections of L6b neurons to L5 might be of particular importance for brain state control. L6b neurons furthermore project to cortical layer 1 (L1) (Clancy and Cauller, 1999) and the hippocampus (Ben-Simon et al., 2022), suggesting an additional role in higher order cognitive functions. L6b is therefore uniquely placed to integrate information from both subcortical and cortical regions, convey signals across L1, L5, and higher order thalamus, and promote cortical activation.

Another important property of L6b neurons is their responsiveness to orexin (also known as hypocretin), a powerful wake-promoting neurotransmitter (De Lecea et al., 1998; Sakurai et al., 1998; Bayer et al., 2004; Wenger Combremont et al., 2016a, 2016b; Zolnik et al., 2024; Messore et al., 2025). Experimental studies have shown that optogenetic activation of orexin neurons promotes awakening (Adamantidis et al., 2007), while optogenetic inhibition promotes sleep (Tsunematsu et al., 2011). The significance of orexin in brain state control is especially apparent in narcolepsy, wherein loss of orexin signalling results in state instability, including sleep attacks and fragmented sleep (Sakurai, 2007). The established view is that orexinergic effects on brain state control are mediated by subcortical circuitry (Sakurai, 2007). However, based on orexinergic projections to the prefrontal cortex and orexinergic induction of spikes in prefrontal cortex slice preparations from rats, it was speculated that the cerebral cortex may contribute to orexin-mediated increases in attention and arousal (Lambe and Aghajanian, 2003). Subsequent work in rat brain slices established the responsiveness of L6b to orexin (and nicotine), and the ability of L6b to potentiate thalamocortical arousal leading to the suggestion that L6b is an essential part of an orexin-gated feed-forward loop driving brain state control (Hay et al., 2015; Zolnik et al., 2026). In support of this, recent findings from our lab with multielectrode array recordings of mouse brain slices show that cortical activation in prefrontal cortex by orexin is impaired in L6b silenced mice (Messore et al., 2025).

We targeted L6b neurons by using the dopamine receptor 1a (Drd1a)-Cre expressing mouse strain Tg(Drd1-Cre)FK164GSat. In this mouse line, Cre is relatively selectively expressed in the lower layers of the cortex (Hoerder-Suabedissen et al., 2018; Zolnik et al., 2024). Subcortically, sparse expression is found in the hippocampus and the striatum, with denser expression in several midbrain nuclei and in the cerebellum (Hoerder-Suabedissen et al., 2018). Drd1a-Cre cortical expression is specific to excitatory neurons and has been quantified to include 25-40% of L6b neurons in the somatosensory cortex (Hoerder-Suabedissen et al., 2018; Zolnik et al., 2020). These Drd1a-Cre neurons in the primary somatosensory cortex receive predominantly long-range intracortical input (Zolnik et al., 2020), project cortically to L1 and L5, and project subcortically selectively to higher order thalamic nuclei (e.g. lateral posterior (LP) and posterior (Po) nuclei), avoiding first order thalamic nuclei (including the dorsal lateral geniculate nucleus, ventrobasal nucleus, and medial geniculate nucleus) (Hoerder-Suabedissen et al., 2018). Drd1a-Cre L6b projections do not exhibit collaterals in the thalamic reticular nucleus (TRN) and form small boutons at their targets (Hoerder-Suabedissen et al., 2018; Casas-Torremocha et al., 2022).

In this study, we continuously recorded EEG/EMG signals across sleep-wake states in “L6b silenced” mice and Cre-negative littermate controls. We achieved L6b silencing by selectively ablating synaptosomal protein of 25 kDa (SNAP25) from dopamine receptor 1a (Drd1a)-Cre expressing neurons in L6b in mice from birth (Hoerder-Suabedissen et al., 2018), a population which has been demonstrated to be orexin-sensitive (Zolnik et al., 2024). The same mouse model was used in previous studies, one of which showed that chronic L6b silencing eventually elicits neurodegeneration (Hoerder-Suabedissen et al., 2019), and another of which indicated that the orexin receptor agonist YNT-185 has differential effects on high-density planar multielectrode array recordings in L6b silenced compared to control animals (Messore et al., 2025); both of these studies indirectly confirm effective silencing. We found that L6b silenced mice show a slowing of theta-frequency oscillations during both wakefulness and REM sleep, and an overall decrease in EEG power centred within the spindle frequency range during NREM sleep. Furthermore, during sleep deprivation, slow EEG power increased to a lesser extent in L6b silenced mice, as compared to control animals. In response to intracerebroventricular (ICV) administration of orexin A and B, both genotypes showed a strong increase in wakefulness, with a minor difference between genotypes wherein orexin A infusion resulted in a greater increase in wake episode duration in L6b silenced compared to control animals, yet it was followed by reduced sleep slow-wave activity. Taken together, our findings suggest that L6b plays an important role in regulating state-dependent cortical dynamics and has a potential role in wake maintenance.

## Results

### The amount of time spent in sleep-wake states is not altered in L6b-silenced mice

In the cortex of L6b silenced mice, Drd1a-Cre expression is most prominent in L6b yet also observed in L6a and occasionally in L5 across cortical regions (Fig. 1, Suppl. T1-T2). No Drd1a-Cre expression was detected in the suprachiasmatic nucleus (Fig. S1).

**Figure 1.**
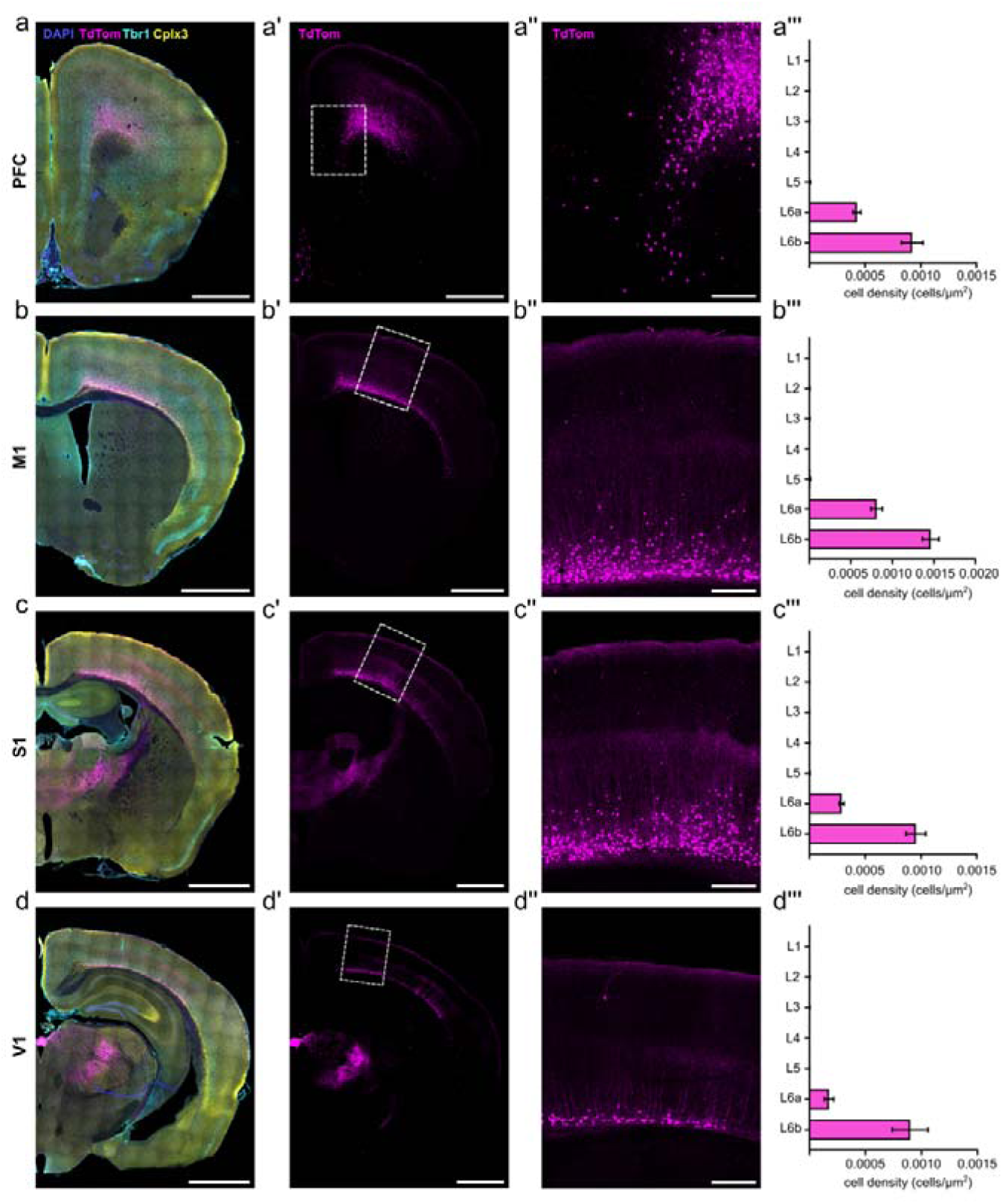
Drd1a Cre positive cells are preferentially localized in L6b across the entire cortical mantle. Drd1a (TdTom) distributions across prefrontal cortex (a-a’’), primary motor cortex (b-b’’), primary somatosensory cortex (c-c’’) and primary visual cortex (d-d’’) in a single example animal, with cell densities averaged across multiple animals (a’’’-d’’’). (a,b,c,d) Hemisection for reference, with DAPI staining for tissue structure (blue), Drd1a-Cre+ cells identified by TdTom expression (magenta). Tbr1 immunostaining (cyan) labels layer 6 and is used to identify the layer 5-6 boundary. Cpxl3 immunostaining (yellow) labels L6b and is used to distinguish L6a and L6b. Tiled images. (a’,b’,c’,d’) Hemisection for reference with only the Drd1a-TdTom channel shown. White boxes indicate the cortical region of interest enlarged in (a’’,b’’,’c’’,d’’). Tiled images. (a’’,b’’,c’’,d’’) Magnified image of the cortical region of interest. (a’’’,b’’’,c’’’,d’’’) Laminar cell densities of Drd1a-Cre;TdTom positive cells. Within-region between-layers comparisons showed that greatest densities are found in L6b, followed by L6a, with sparse to no Drd1a expression in upper layers (Cortical layer, F_(6,112)_=319.6, p<0.0001) in each cortical region analysed. Please refer to Supplemental table T1 and T2 for exact combinatoric inter-layer and inter-region test results, respectively. Data represented as mean ± SEM. Scale bar a-d and a’-d’, 1000 µm. Scale bar a’’-d’’, 200 µm. Experimental replicates, PFC (n=5), M1 (n=6), S1 (n=6), V1 (n=3). For each animal, 3 technical replicates were used. Images were obtained with a spinning-disk confocal microscope.

First, we performed EEG and EMG recordings in control and L6b silenced mice over an undisturbed baseline recording using an approach established in our previous work (Yamagata et al., 2021). Electrodes were implanted above the frontal and the occipital cortex and referenced against the cerebellum to monitor state-specific brain oscillations including EEG slow-wave activity (SWA, 0.5-4Hz) and spindle-frequency (10-15 Hz) activity, which are both more prominent in anterior derivations in mice (Vyazovskiy et al., 2004; Vyazovskiy and Tobler, 2005a; Blanco-Duque et al., 2024), and theta-activity, which is typically more pronounced in posterior electrodes (Huber et al., 2000; McKillop et al., 2021) (Fig. 2A). Visual inspection of EEG traces confirmed that the key features of vigilance states were clearly identifiable in both genotypes (Fig. 2B). We then analysed the daily profiles of SWA and spectrograms (representative individuals: Fig. 2C), which revealed the typical spectral signatures of waking, NREM, and REM sleep in both control and L6b silenced animals. Specifically, we observed that high SWA (0.5-4 Hz) was prominent during epochs scored as NREM sleep, while theta activity (4-10 Hz) was strongest during epochs scored as REM sleep. Plotting individual hypnograms indicated that, as expected in all animals regardless of genotype, sleep occurred mostly during the light phase and wakefulness was more consolidated during the dark period as expected for mice in laboratory conditions (Fig. 2D). This analysis reassured us that functionally silencing neocortical L6b neurons does not result in the emergence of any atypical brain oscillations or states under baseline conditions which may contaminate spectral analyses or compromise our ability to annotate vigilance states using conventional criteria.

**Figure 2.**
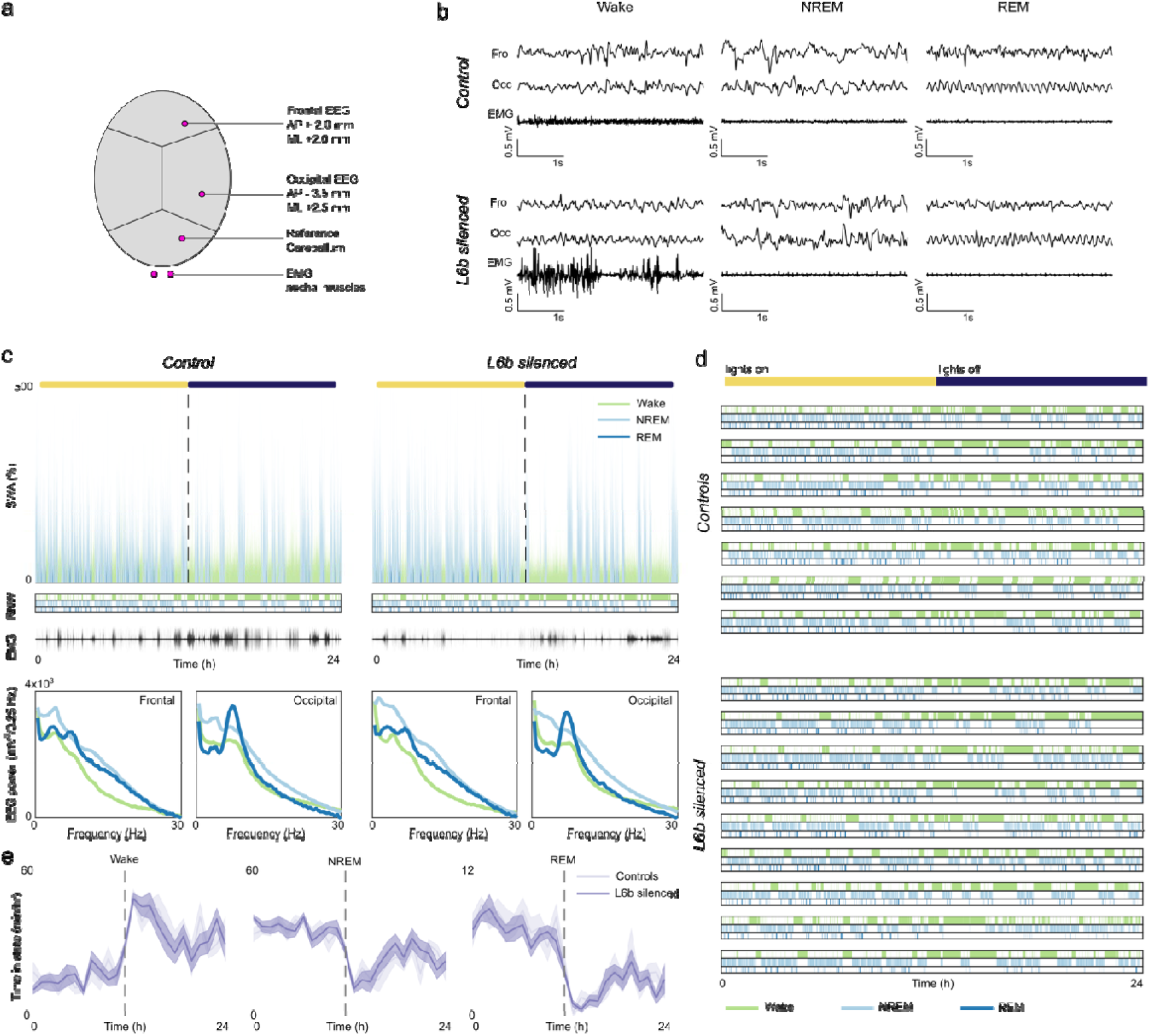
Daily sleep-wake architecture is unchanged in L6b silenced animals. a. Position of electrodes. The frontal and occipital EEG (expressed as mm distance from Bregma in midline (ML) and anteroposterior (AP) directions) were referenced against a cerebellar screw. Two EMGs were implanted in the nuchal musculature and referenced against one other. b. Representative frontal EEG, occipital EEG and EMG traces during the respective vigilance states in a L6b silenced and a control animal show similar patterns of activity, allowing blinded scoring of vigilance states. c. 24-hour profiles of slow wave activity (SWA) in the frontal EEG (% of mean), EMG activity (arbitrary units), and vigilance-specific spectrograms in a representative L6b silenced animal and control animal. The bar on top represents the duration of the light phase (yellow) and dark phase (dark blue). d. Hypnograms for all individual animals. e. Daily time course of wakefulness, NREM and REM sleep were comparable in L6b silenced and control animals. Controls n=7, L6b silenced n=9.

Across 24 hours, L6b silenced and control animals spent comparable time in wakefulness (controls 41.8±1.01% vs L6b silenced 40.9±1.91%, t_(14)_=0.357, p=0.726), NREM sleep (controls 50.2±1.01% vs L6b silenced 51.5±1.83%, t_(14)_=0.8672, p=0.554), and REM sleep (controls 8.05±0.695% vs L6b silenced 7.52±0.394%, t_(14)_=0.710, p=0.491). In addition, the time course of vigilance states across 24h did not reveal significant differences in the distribution of vigilance states across the light-dark cycle between genotypes (Fig. 2E).

Next, we addressed whether L6b silencing affects continuity of sleep-wake states. We found that the average duration of individual episodes of wakefulness (controls 14.3±1.29 min vs L6b silenced 14.7±1.43 min, t_(14)_=0.1663, p=0.8703), NREM sleep (controls 4.28±0.200 min vs L6b silenced 4.50±0.298 min, t_(14)_=0.5671, p=0.5796), and REM sleep episodes (controls 1.04±0.0344 min vs L6b silenced 1.10±0.0523 min, t_(14)_=0.9792, p=0.3441) did not differ between genotypes. Likewise, the number of brief awakenings — short regular interruptions of sleep by epochs resembling wakefulness, characterised by EEG activation and an increase in EMG signal lasting ≤16 s (Hauglund et al., 2025) —was not altered (controls 51.1±2.04 h^-1^ vs L6b silenced 51.6±3.97h^-1^, t_(12)_=0.1205, p=0.9061).

Consistent with EEG/EMG defined vigilance states, passive infrared monitoring of locomotor activity, undertaken in a separate circadian phenotyping experiment, revealed only minor differences between genotypes. In light-dark conditions, the endogenous period was close to 24h in both genotypes (control 24.0 ± 0.006 h, L6b silenced 24.0 ± 0.01 h, t_(11)_=0.1422, p=0.8895); however, the active phase, alpha, was slightly longer in L6b silenced animals (control 12.5 ± 0.151 h vs L6b silenced 13.4 ± 0.348 h, t_(11)_=2.443, p=0.0326). The number of activity bouts per day, the duration of activity bouts, and the number of activity counts per bout did not differ with L6b silencing. Constant darkness experiments revealed that neither the intrinsic period nor alpha differed between genotypes (period: control 23.8 ± 0.0605 h vs L6b silenced 23.6 ± 0.0626 h, t_(11)_=1.721, p=0.1133; alpha: control 15.3 ± 1.46 h vs L6b silenced 14.9 ± 0.737 h, t_(11)_=0.2313, p=0.8213). The number of activity bouts per day, the duration of individual bouts, and the number of activity counts per bout were similar between genotypes, but the number of activity counts per day showed a trend towards lower activity in L6b silenced animals (t_(11)_=2.045, p=0.0656).

### Layer 6b silencing leads to state-dependent changes in the EEG

After examining daily sleep-wake amount, time course, and architecture, we investigated the effect of L6b silencing on EEG power spectra. During wakefulness, only minor changes were observed in the frontal derivation (Fig. 3A). However, differences were more pronounced in the occipital EEG, especially in the higher theta frequency range where power was significantly reduced due to a leftward shift of theta peak frequency (theta peak frequency during wake: controls 7.42±0.300 Hz vs L6b silenced 5.78 ± 0.485 Hz, Welch’s t test, t_(12.40)_=2.873, p=0.0136) (Fig. 3A, 3D, 3E). Higher frequencies were not significantly affected (Fig. S2, Suppl. T3). During REM sleep, L6b silencing resulted in a reduction of frontal EEG power in the theta and higher frequency ranges (Fig. 3A, Fig. S2, Suppl. T3). In the occipital derivation, we observed a leftward shift of theta-peak frequency in L6b silenced animals (Fig 3D, 3E) (during REM: controls 7.58±0.0833 Hz vs L6b silenced 7.11 ± 0.0735 Hz, Welch’s t test, t_(11.47)_=4.250, p=0.0012). During NREM sleep, L6b silencing markedly reduced frontal EEG spectral density across nearly the entire frequency range examined, from 3Hz upward (Fig. 3A, Fig S2, Suppl. T3). No differences in the occipital EEG were apparent across low frequencies, while a slight decrease in frequencies above 80 Hz was observed (Fig. S2). To further investigate state-specificity of these differences, we examined changes in EEG spectra around NREM-REM sleep transitions (Fig. 3B). This revealed that the EEG power decrease in NREM sleep was especially prominent in the frontal derivation around the spindle-frequency range (Blanco-Duque et al., 2024), while a shift in theta-peak frequency during REM sleep was especially apparent in the occipital derivation (Fig. 3E). Importantly, the reduced EEG power during NREM sleep was not specific to NREM sleep immediately before NREM-REM state transitions, as normalising EEG spectra in the 32 seconds of NREM sleep preceding NREM-REM transitions to overall NREM sleep spectral power density on baseline day abolished differences between genotypes (Fig. 3C).

**Figure 3.**
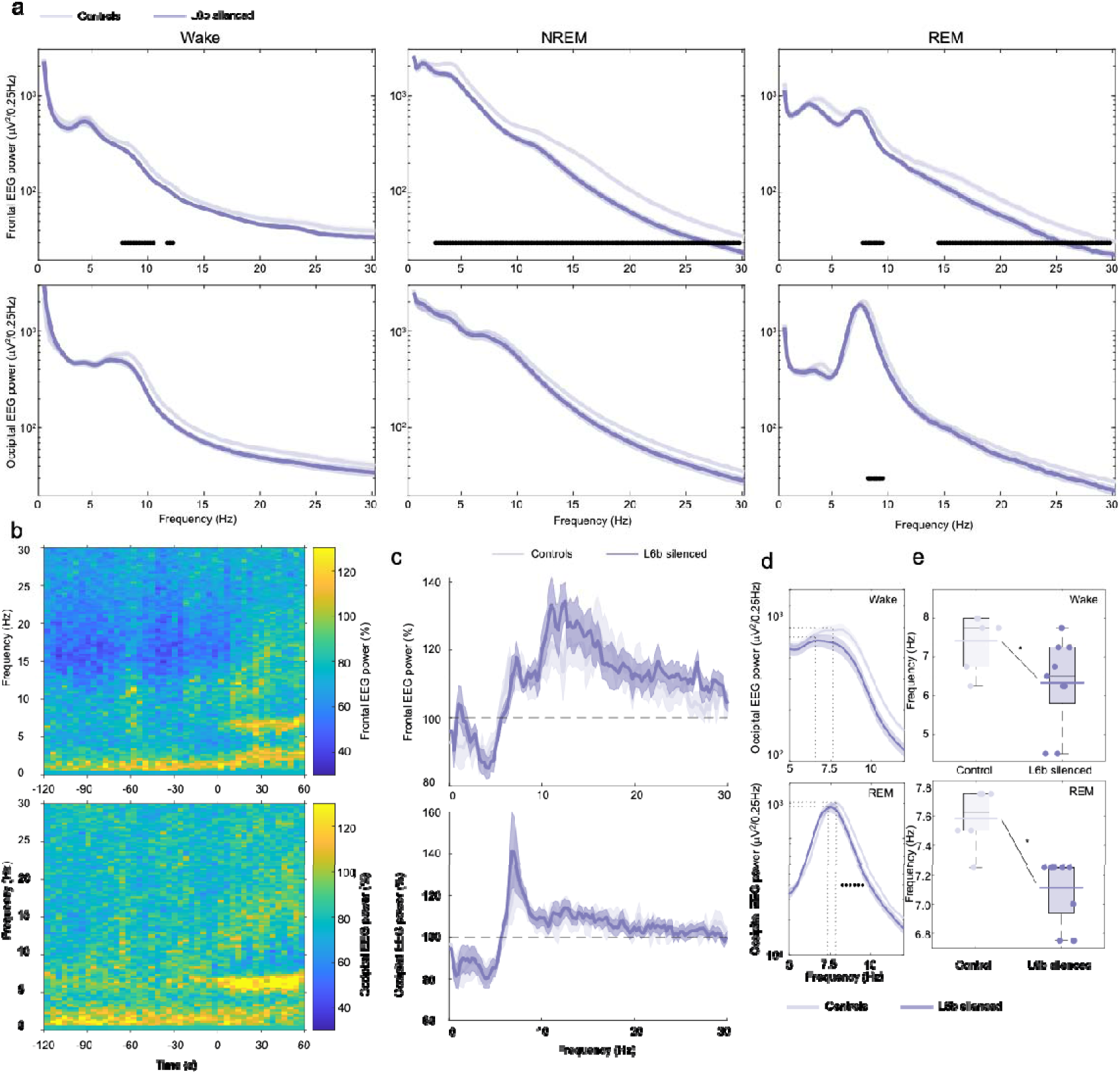
Vigilance state specific EEG spectra are changed in L6b silenced animals. a. EEG spectral power in the frontal and occipital derivation during Wake, NREM and REM sleep in L6b silenced and control animals. Filled dots show bins with significant differences between genotype groups after comparison with post hoc tests in 0.25 Hz bins when there was a significant genotype x frequency interaction with two-way ANOVA. Frontal EEG, Controls n=7, L6b silenced n=9. Occipital EEG, Controls n=6, L6b silenced n=9. b. EEG spectral power in the 2 minutes preceding and 1 minute following NREM-REM transitions averaged across all NREM-REM transitions across 24 hours, with average power in L6b silenced animals relative to control animals in percentages. Top, frontal EEG, controls n=6, L6b silenced n=9. Bottom, occipital EEG, controls n=6, L6b silenced n=9. c. EEG spectral power during NREM sleep in the 32 seconds preceding the NREM-REM transition relative to the EEG power in NREM sleep across 24 hours, in the frontal (top, controls n=7, L6b silenced n=9) and occipital (bottom, controls n=6, L6b silenced n=9) EEG. d. Enlarged representation of the occipital EEG spectral power shown in (a) during wakefulness (top) and REM sleep (bottom). Filled dots mark significant genotype differences in 0.25-Hz bins with post hoc tests when there was significant genotype x frequency bin interaction in the two-way ANOVAs. Controls n=6, L6b silenced n=9. Dotted lines illustrate the EEG spectral power and frequency of the theta peak. e. Peak theta frequency in the occipital EEG during wakefulness (top) and REM sleep (bottom) for control and L6b silenced animals. Asterisks mark significant differences between genotypes. Controls n=6, L6b silenced n=9.

### Layer 6b silencing affects spectral signatures of sleep pressure during waking and sleep

Next, we investigated the effects of L6b silencing on homeostatic sleep regulation by undertaking 6 hours of sleep deprivation (SD) starting from light onset. This manipulation is usually associated with an increase in EEG indices of cortical arousal (Vyazovskiy and Tobler, 2005b; Vassalli and Franken, 2017; Yamagata et al., 2021), intrusion of sleep-like patterns of activity in wake EEG (Vyazovskiy et al., 2011), and subsequently increased EEG SWA during NREM sleep (Thomas et al., 2020). SD was successful in all animals, with only minimal sleep observed during the 6-hour SD procedure (controls 15.0±3.73 min vs L6b silenced 14.3±3.97 min, t_(14)_=0.1240, p=0.9030). Animals fell asleep soon after the end of SD, where the latency to the first consolidated NREM sleep did not differ significantly between genotypes (controls, 6.75±2.92 min, L6b silenced 11.2±2.56 min, t_(14)_=1.153, p=0.2680). Representative hypnograms of two individual animals during the sleep deprivation experiment are shown in Fig 4A.

**Figure 4.**
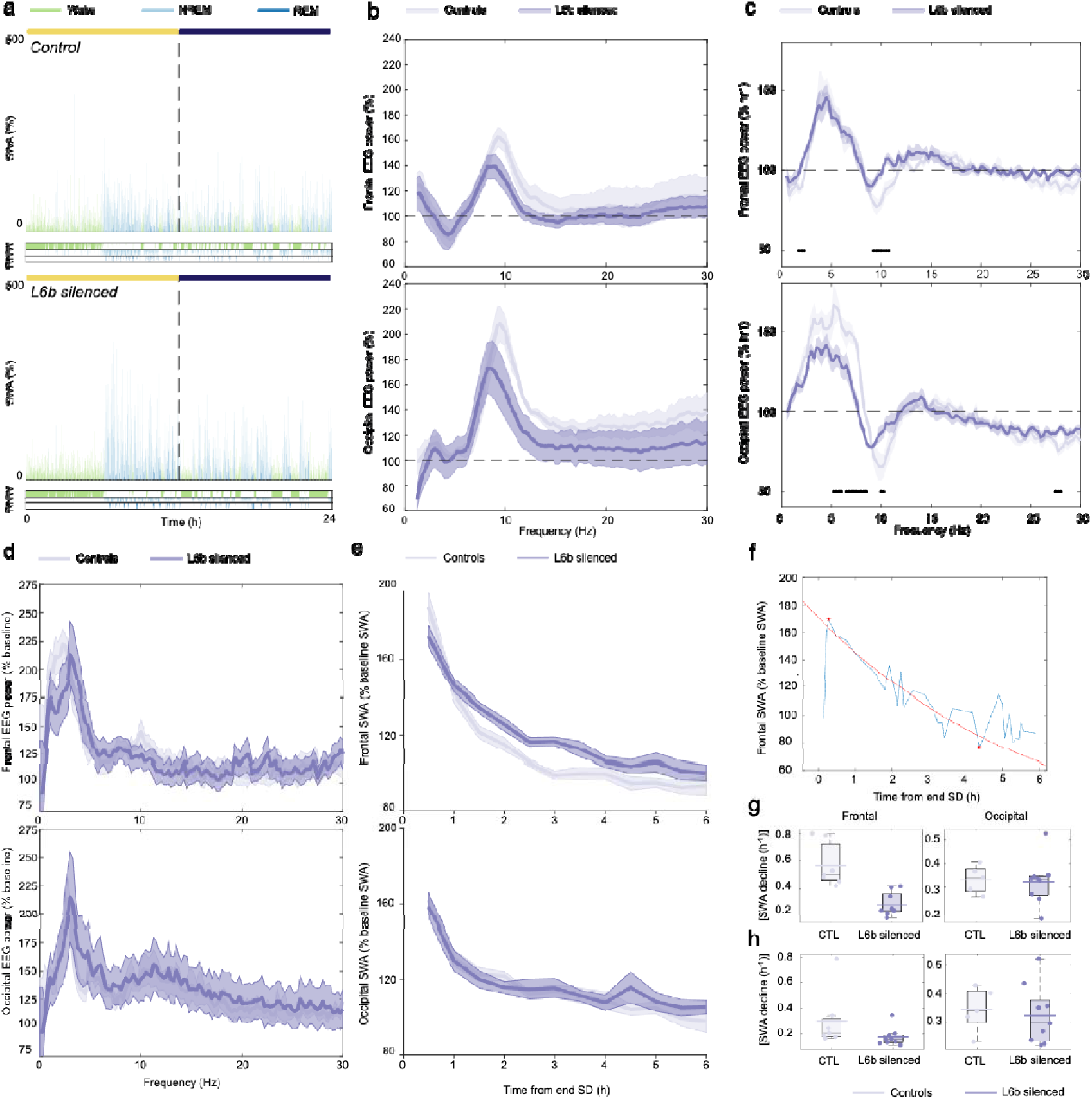
The response to sleep deprivation is altered in L6b silenced animals. a. 24-hour profiles of SWA (0.5-4.0 Hz) after sleep deprivation in a representative control and L6b silenced animal, with hypnograms plotted below. The bar on top marks the duration of the light phase (yellow) and dark phase (dark blue). b. EEG power during sleep deprivation in L6b silenced and control animals, normalised to wakefulness spectra on baseline day. c. EEG spectral power in the sixth (final) hour of sleep deprivation normalised to the first hour of sleep deprivation. Filled circles represent 0.25 Hz bins where post hoc tests showed a significant genotype difference, when two-way ANOVA showed a significant genotype x frequency interaction. d. Spectral power in the frontal EEG during the first 30 minutes of NREM sleep following sleep deprivation. e. Time course of SWA (0.5-4.0 Hz) during NREM sleep across the six hours following sleep deprivation. f. The rate of SWA decline was approximated with an exponential fit, the method is shown for the frontal EEG from a representative individual animal. g. Absolute exponent coefficients for an exponential fit across only the first two hours following sleep deprivation. h. Absolute exponent coefficients for an exponential fit across only the first six hours following sleep deprivation. Frontal EEG, controls n=7, L6b silenced n=9. Occipital EEG, controls n=6, L6b silenced n=8.

First, we analysed wake EEG spectra during sleep deprivation. In both control and L6b silenced animals, a broadband increase in EEG power, most prominently in the theta frequency range, was observed during sleep deprivation compared to undisturbed wakefulness (Fig. 4B; see Suppl. Table T4-T7 for a statistical comparison per genotype between normalised spectra and a theoretical mean=1 by multiple one-sample t tests). This may indicate a heightened wake “intensity” or level of arousal (Krone et al., 2021; Yamagata et al., 2021) induced by continuous provision of novel objects. Interestingly, when we evaluated the change in wake EEG spectral power from the first to the last hour of sleep deprivation, L6b-silenced animals showed an attenuated increase in spectral power density in the 9.5–11 Hz and 5.5-9.0 Hz range compared to controls, in the frontal and occipital derivation, respectively (Frontal: Frequency x Genotype, F(118,1652)=1.597, p<0.0001, post-hoc test significant 9.5-11.0 Hz; Occipital: Frequency x Genotype, F_(118,1298)_=4.543, p<0.0001, post hoc test significant 5.5-9.0 Hz)(Fig. 4C).

In both genotypes, SD was followed by an initial increase in NREM sleep EEG power in the slow-wave frequency range (Fig. 4D), with an expectedly smaller change in the occipital derivation compared to the frontal derivation. Plotting the time course of NREM sleep SWA across the first 6 h after SD revealed a typical dynamic with higher values of SWA at the beginning of recovery sleep followed by its progressive decline in both genotypes and in both derivations (Fig. 4E). An exponential function was fitted to the time course of NREM sleep SWA dissipation after the end of SD to estimate the time constant in L6b silenced animals and controls (Fig. 4F). While the optimal timeframe for this analysis is difficult to define, the most prominent effects of layer 6b silencing were observed early after the end of SD. Specifically, when the time constant was estimated across the first 2 hours after the end of SD only, it was significantly lower in L6b silenced animals compared to controls in the frontal EEG (controls -0.57 ± 0.060 h-1 vs L6b silenced -0.28 ± 0.029 h-1, t_(8.723)_=4.286, p=0.0022, unpaired Welch’s t test)(Fig. 4G). SWA dissipation in the occipital EEG did not differ between genotypes (controls -0.33 ± 0.026 h-1 vs L6b silenced -0.34 ± 0.029 h-1, t_(10.73)_=0.2732, p=0.79). However, when the time frame was expanded to 6 hours after the end of SD, there was no significant difference between genotypes in the frontal EEG (controls –0.30 ± 0.084 h-1 vs L6b silenced –0.17 ± 0.023 h-1; t_(6.925)_= 1.451, p= 0.19, unpaired Welch’s t test) or occipital EEG (controls –0.12 ± 0.017 h-1 vs L6b silenced –0.12 ± 0.018 h-1, t_(10.46)_= 0.1675, p=0.87) (Fig. 4H).

### Orexin promotes wakefulness in L6b silenced and control animals

Subsequently, we compared effects of intracerebroventricular (ICV) orexin administration between control and L6b silenced animals. Histology confirmed that cannulas were positioned in the right lateral ventricle in all animals. Vehicle, orexin A, or orexin B were infused at light onset (Fig. 5A,B). Visual inspection of EEG traces did not reveal any abnormalities or obvious differences between genotypes (Fig. S3), therefore sleep scoring was performed blindly using established criteria (Yamagata et al., 2021). No abnormal oscillatory activities were revealed by subsequent spectral analysis in either genotype or derivation (representative individual examples: Fig. S3).

**Figure 5.**
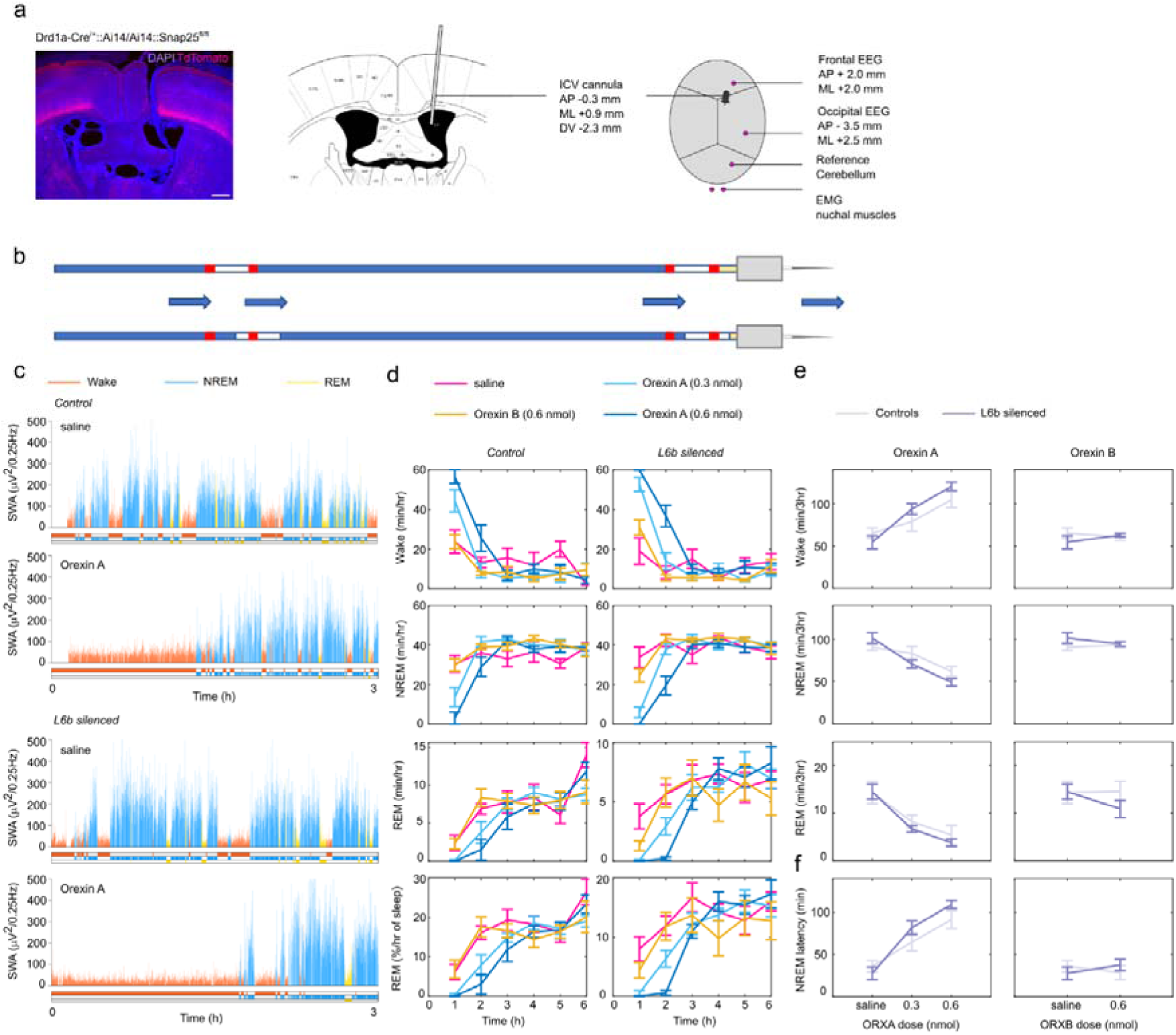
Intracerebroventricular infusion of orexin A promotes wakefulness in L6b silenced and control animals. a. Histological and schematic overview of the intracerebroventricular infusion canula and EEG/EMG implant, with position of the cannula and electrodes defined as coordinates from bregma in the anterioposterior (AP) direction and midline (ML) direction, and cannula dorsoventral position (DV) from the surface of the dura. b. Schematic overview of the infusion procedure with tubing front-filled with orexin or vehicle solution and backfilled with saline. c. Time course of SWA in the 3 hours following infusion of vehicle and 0.6 nmol orexin A in a representative control and L6b silenced animal. d. Time course of vigilance states in the first 6 hours after infusion of vehicle (saline), a lower dose of orexin A (0.3 nmol), a higher dose of orexin A (0.6 nmol), orexin B (0.6 nmol) in L6b silenced and control animals. Controls n=4, L6b silenced n=7. e. Total amount of wake, NREM and REM sleep in the first 3 hours after infusion of orexin A or orexin B. Controls n=4, L6b silenced n=7. f. The latency to consolidated NREM sleep was increased after infusion of orexin A (left column) but not ORXB (right column). Controls n=4, L6b silenced n=7.

Sleep architecture was considerably affected by orexin administration. As depicted in individual examples, while vehicle administration had no effect on vigilance states, orexin induced prolonged wakefulness in both genotypes (Fig. 5C, Fig. S4). The 6-hour time course of vigilance states revealed a pronounced increase in wakefulness and decrease in sleep after both doses of orexin A in both genotypes, while effects of orexin B were less prominent (Fig. 5D). Based on these findings that illustrate that the effects dissipated beyond three hours (Fig. 5D) and reports in literature regarding the duration of action of orexin (Piper et al., 2000; Huang et al., 2001; Mieda et al., 2011), the time frame used to analyse sleep architecture was defined as the time from the end of the infusion to three hours post-infusion. During this interval, ORXA administration markedly and dose-dependently increased the time spent in wakefulness (Fig. 5E; ORXA Dose, F_(1.316,11.18)_=27.56; p=0.0001); there was no difference between genotypes in the total amount of time spent in wakefulness during the time frame analysed (ORXA Dose x Genotype, F_(2,17)_=2.597, p=0.10), even though the mean duration of individual wake episodes within that time frame was increased to a greater extent in L6b silenced animals (ORXA Dose x Genotype, F_(2,22)_=4.376, p=0.0251). The amount of NREM sleep and REM sleep decreased with ORXA administration (NREM sleep: Vehicle controls 91.1 ± 4.73 min vs L6b silenced 96.3 ± 4.61 min, ORXA controls 65.4 ± 6.08 min vs L6b silenced 51.1 ± 4.43 min, ORXA Dose F_(1.629,14.66)_=22.30, p<0.0001; REM sleep: Vehicle controls 15.3 ± 2.75 min vs L6b silenced 14.8 ± 1.78 min, ORXA controls 6.23 ± 2.39 min vs L6b silenced 4.07 ± 0.818 min, ORXA Dose, F_(1.057,9.516)_=21.23, p=0.001). In contrast, ORXB did not have a significant effect on vigilance states (as compared to vehicle: Wake, ORXB Dose F_(1,9)_=0.02874, p=0.8691; NREM sleep, ORXB Dose, F_(1,9)_=0.00898, p=0.9266; REM sleep, ORXB Dose, F_(1,20)_=1.193, p=0.2878).

The latency to onset of consolidated NREM sleep significantly increased following ORXA infusion in both genotypes (Vehicle: controls 36.2 ± 6.94 min vs L6b silenced 27.7 ± 7.58 min, ORXA: controls 91.4 ± 10.5 min vs L6b silenced 109 ± 4.82 min, ORXA Dose, F_(2,22)_=50.10, p<0.0001; ORXA Dose x Genotype, F_(2,22)_=2.376, p=0.1164))(Fig. 5F), as did the average duration of wake episodes (Vehicle: controls 19.8 ± 4.92 min vs L6b silenced 10.6 ± 1.40 min; ORXA: controls 25.8 ± 5.39 min vs L6b silenced 39.7 ± 7.53 min, ORXA Dose, F_(2,22)_=8.995, p=0.0014). Lastly, both ORXA and ORXB infusion significantly increased the number of brief awakenings from NREM sleep in both genotypes (Vehicle: controls 69.59 ± 2.19 hr^-1^, L6b silenced 55.04 ± 2.84 hr^-1^, ORXA: controls 83.14 ± 22.69 hr^-1^, L6b silenced 93.36 ± 2.68 hr^-1^, ORXB: controls 85.53 ± 14.80 hr^-1^, L6b silenced 71.91 ± 7.02 hr^-1^; ORXA Dose, F_(1.460,8.759)_=7.907, p=0.0149, Mixed-effects model; ORXB Dose, F_(1,7)_=6.431, p=0.0389, two-way ANOVA). Overall, these results confirm our prediction that ORXA promotes wakefulness.

### Orexin-induced wakefulness leads to an increase in homeostatic sleep pressure

Since orexin promotes wakefulness, its administration at light onset can be regarded as a form of pharmacological sleep deprivation. While ORXA administration significantly increased the time spent in wakefulness in the first 3 hours, wakefulness returned to normal levels from the third hour onwards and 24-hour percentages in respective vigilance states were comparable after ORXA and vehicle infusions independent of genotype (ORXA dose effect: Wake, F_(1.261,13.87)_=1.431, p=0.26; NREM sleep, F_(1.270,13.97)_=1.615, p=0.23; REM sleep, F_(1.559,17.15)_=2.042, p=0.17. ORXA dose x genotype interaction: Wake, F_(2.22)_=1.247, p=0.31; NREM sleep, F_(2,22)_=1.415, p=0.26; REM sleep, F_(2,22)_=0.1588, p=0.85) (Fig. S5A). The 24-hour percentage of time spent in REM sleep was reduced in L6b silenced animals compared to controls in both vehicle and orexin conditions without a treatment x genotype interaction (Vehicle: controls 9.46 ± 0.683% vs L6b silenced 7.90 ± 0.379%; ORXA: controls 8.67 ± 0.373% vs L6b silenced 7.44 ± 0.257%; ORXB: controls 8.40 ± 0.313% vs L6b silenced 7.30 ± 0.710%; ORXA, Genotype effect, F_(1,11)_=8.618, p=0.014, Genotype x ORXA dose interaction, F_(2,22)_=0.1588, p=0.85; ORXB, Genotype effect, F_(1,11)_=7.130, p=0.022), Genotype x ORXB interaction, F_(1,11)_=0.1194, p=0.74) (Fig. S5A, Fig. S5B). Interestingly, while SWA during the initial 3 hours after sleep onset following ORXA administration (Fig. S5C) declined at a similar rate in both genotypes (Frontal, Time x Genotype F_(5,55)_=0.4621, p=0.8027, Occipital, Time x Genotype, F_(5,55)_=1.373, p=0.2485), the overall levels of SWA across the first 3 hours of sleep were lower in the occipital derivation in L6b silenced animals compared to controls (Occipital EEG, Genotype effect, F_(1,11)_=6.818, p=0.0242).

## Discussion

### L6b plays a role in regulation of arousal states

In this study, we examined sleep-wake regulation in the “L6b silenced” mouse model, wherein the synaptic protein Snap25 is chronically ablated from birth in the Drd1a-Cre expressing subpopulation of L6b (Drd1a-Cre; Snap25^fl/fl^;Ai14) across the entire cortical mantle. The distribution of vigilance states, and measures of sleep consolidation and sleep fragmentation did not differ between L6b silenced and control animals at 12 weeks of age. However, during wakefulness, EEG activity in the theta-frequency range, the signature of active or exploratory wakefulness (Vyazovskiy et al., 2006; Vassalli and Franken, 2017), was characterised by a lower peak frequency in L6b-silenced animals, while in REM sleep there was a reduction in theta peak frequency, along with a reduction in gamma-frequency power. We posit that silencing L6b impairs the establishment of an activated brain state, which is required to generate a state of arousal and alertness.

Our findings are in line with previous *in vitro* studies showing that L6b neurons can be activated by orexin, neurotensin, noradrenaline, histamine, and dopamine – all of which are powerful promoters of arousal (Bayer et al., 2004; Wenger Combremont et al., 2016a, 2016b). Notably, data from Zolnik *et al*. show that optogenetically activating the same Drd1a-Cre+ subpopulation of L6b neurons in head-restrained awake mice can promote conversion from delta oscillations to gamma oscillations, thus promoting a state of cortical activation resembling wakefulness and REM sleep (Zolnik et al., 2024). Our study of long-term recordings in freely behaving mice supports this same conclusion; L6b is involved in generating or maintaining active substates of wakefulness.

Anatomically, L6b is ideally positioned to contribute to the regulation of levels of arousal and vigilance, which are known to exhibit a rich dynamic in both waking and sleep (Andrillon, 2023). L6b receives input predominantly from long-range intracortical projections and projects selectively to higher order thalamic nuclei (Hoerder-Suabedissen et al., 2018; Zolnik et al., 2020, 2024). Therefore, L6b can integrate inputs from wake-promoting subcortical neuromodulatory areas, as well as information from distant cortical areas and translate this into wide-spread cortical activation with additional relay via higher-order thalamic nuclei. Higher order thalamic nuclei can themselves also be directly activated by orexin (Bayer et al., 2002), and orexinergic activation of L6b can enhance the integration of higher order thalamic inputs in L6a (Hay et al., 2015).

### A role of cortical L6b in brain oscillations

Silencing a subpopulation of L6b not only changed cortical activity during wakefulness and REM sleep, but also during NREM sleep. This was reflected in a reduction in NREM sleep EEG power across a broad frequency range, including delta (0.5-4 Hz), theta (5-10 Hz), sigma (10-15 Hz), and gamma (30-100 Hz) frequencies. Slow wave activity (SWA, 0.5-4 Hz) including slow waves and delta waves is a marker of sleep intensity and reflects the levels of sleep pressure, which builds up as a function of preceding wakefulness duration (Borbély, 1982; Achermann and Borbély, 2003). The reduction of delta power during NREM sleep in L6b silenced animals agrees with the reduction of theta power during wakefulness, since the theta substate of wakefulness drives the build-up of sleep pressure (Vyazovskiy and Tobler, 2005b; Vassalli and Franken, 2017; Yamagata et al., 2021). EEG activity in the delta frequency band is thought to reflect spatial and temporal synchronization across large neuronal populations undergoing up- and down states (Steriade et al., 1993; Vyazovskiy et al., 2009; Thomas et al., 2020), therefore the reduction in SWA in L6b silenced animals may suggest a role for L6b in network synchrony.

Sigma power (10-15 Hz) represents the frequency range of sleep spindles (Blanco-Duque et al., 2024), which are waxing and waning bursts of activity that are thought to originate from interactions between cortex, thalamus, and the thalamic reticular nucleus (TRN) (Steriade et al., 1993; Crunelli et al., 2006). Sleep spindles have been linked to memory consolidation in sleep and are posited to facilitate the transfer of information to long-term storage (Helfrich et al., 2018; Hahn et al., 2019). Thus, the reduction in sigma power in L6b silenced animals suggests that L6b may be involved in these functions. L6b does not directly project to the TRN (Hoerder-Suabedissen et al., 2018), yet there are projections from L6b to L5 (Zolnik et al., 2024), and subpopulations of L5 innervate segments of the TRN in an area-specific pattern (Carroll et al., 2022; Hádinger et al., 2023), therefore L6b may indirectly influence sleep spindle generation via L5.

In addition, the EEG peak theta frequency was slower in L6b-silenced animals. A similar reduction was observed as a result of a global deletion of the GluA1 subunit of AMPA receptors (Ang et al., 2018) and in animals with a SNAP25-ablation in L5 pyramidal neurons an dentate gyrus granule cells (Krone et al., 2021). While a relatively simple explanation, such as a modest reduction in brain temperature (Deboer, 2002), cannot be excluded, the neurophysiological mechanisms underlying this shift in theta peak frequency cannot be reliably inferred from EEG recordings alone. Cortical EEG signals are likely dominated by volume-conducted activity, and therefore resolving the underlying mechanisms would require additional intracortical and intrahippocampal recordings.

### An active role for the neocortex in sleep regulation

We showed recently that state regulation is impaired in mice that have a reduction in network excitability because of a mutation in the synaptic vesicle protein VAMP2 (Guillaumin et al., 2025). These findings are broadly consistent with the view that brain state transitions start at the local level and may or may not converge to a ‘global’ brain state transition (Krueger et al., 2008; Thomas et al., 2020; Andrillon et al., 2024; Guillaumin et al., 2025). In the current study, we showed that SNAP25-ablation in a subpopulation of excitatory L6b neurons across the entire cortical mantle is associated with changes in oscillatory activity during sleep and wakefulness. Mice with a SNAP25-ablation in a subpopulation of cortical L5 across the whole cortical mantle, however, show a substantial reduction in time spent asleep without clearly affecting global EEG oscillations (Krone et al., 2021). These findings suggest that different brain regions or distinct subpopulations of projection neurons in infragranular layers have differential contributions in the dynamics of wake-sleep states or state intensity.

The fact that SNAP25-ablation in both L5 and L6b affects cortical signatures of sleep and wakefulness – one in the time spent in different vigilance states, and the other in state-specific oscillations, respectively – strongly suggests that sleep is not only regulated via subcortical systems forming a ‘sleep wake switch’ at a global level (Saper et al., 2010), but that the neocortex is also actively involved in regulating its own state, likely through both local and distributed intra- and extracortical circuits (Krueger et al., 2008). Recent work has shown that activity in the neocortex is involved in cortical synchrony (Ratliff et al., 2024), regulation of sleep depth and duration (Vaidyanathan et al., 2021), sleep homeostasis (Kon et al., 2024), sleep-preparatory behaviour (Tossell et al., 2023), and REM sleep initiation (Wang et al., 2022; Hong et al., 2023) and progression (Dong et al., 2022), all supporting an active role of the neocortex in brain state regulation. Yet, our current findings also support an important role of subcortical nuclei in the control of local and global cortical states of arousal. In addition to orexinergic effects, this likely extends to other neuromodulatory systems including acetylcholine, noradrenaline, dopamine, neurotensin, serotonin, and histamine (Constantinople and Bruno, 2011; Wenger Combremont et al., 2016b, 2016a; Case et al., 2017; Pal et al., 2018; Parkar et al., 2020; Osorio-Forero et al., 2021; Dean et al., 2022; Bréant et al., 2026).

### Limitations

This study was exploratory in nature from the beginning, and there are several key points that need to be addressed in future studies. First, the Drd1a-Cre driver line FK164 was selected amongst various Drd1a-Cre driver lines that are currently available because of its Cre-expression that is relatively specific to the neocortex. Yet, sparse subcortical expression is still found in this line, in hippocampus, striatum, several midbrain nuclei and in the cerebellum (Hoerder-Suabedissen et al., 2018), and it cannot be excluded that subcortical populations contribute to the effects on brain state control that were detected. Also, within cortex Drd1a-Cre expression is not limited to L6b, with some expression also detected in L6a. Since projection pattern and responsiveness to orexin are mutual across Drd1a-Cre neurons in L6a and L6b, this issue may not be vital, but further characterisation of both populations is needed to solidify the assumption that these cells are part of the same population. In the current study, only male mice were used, because our experimental protocol precluded the possibility of accurately monitoring the oestrous cycle, which has marked effects on brain activity, arousal and vigilance states. We therefore decided to use male mice only for the current study but are planning to use both sexes in future work. Moreover, in the current study, we investigated the effect of chronic manipulation of the Drd1a positive L6b population in freely moving animals. With these findings, we complement previous work on acute manipulation of the same L6b population (Drd1a-Cre) in head-fixed animals (Zolnik et al., 2020, 2024). Data in freely moving animals can aid in understanding brain processes in a more naturalistic setting. Moreover, chronic rather than acute manipulation could offer insights into pathophysiological mechanisms in neurodevelopmental disorders such as schizophrenia, which may be linked to chronic abnormal network development. Even though the current study thus adds valuable longer-term data on chronic manipulations in freely moving animals, next steps should also include acute and spatially restricted manipulations of L6b neurons using a viral vector strategy and intracranial recordings to strengthen insights into the exact cortical area and corticothalamic networks involved. In particular, manipulations restricted to the prefrontal cortex would be of interest due to recent findings point towards a role for this cortical region in arousal from anaesthesia and slow wave sleep (Pal et al., 2018; Parkar et al., 2020; Dean et al., 2022). Finally, the exclusion of some animals from the final data set for technical reasons, limited our ability to draw strong conclusions regarding the effects of orexin on wake EEG spectra, and future studies will be required to address this.

## Methods

### The ‘layer 6b silenced’ mouse model

In this study, the gene encoding the synaptic protein Synaptosomal Associated Protein of 25 kDa (SNAP25), which mediates regulated synaptic vesicle release (Hanson et al., 1997; Poirier et al., 1998; Sutton et al., 1998), was selectively ablated from a subpopulation of L6b neurons to render these functionally ‘silenced’ from the first week of postnatal period. While whole-mouse *Snap25* knockout is impossible due to embryonic lethality (Molnár et al., 2002; Washbourne et al., 2002), embryos with selective *Snap25* elimination from specific cortical projection neurons are viable. *Snap25* ablation from L5, L6a, and L6b, respectively, initially results in normal cortical development, with loss of synapses, demyelination, microglia activation, axonal and neuronal degeneration typically observed at later time points (Hoerder-Suabedissen et al., 2019; Korrell et al., 2019; Vadisiute et al., 2022).

Layer 6b can be targeted in the Tg(Drd1a-cre)FK164Gsat/Mmucd mouse (Drd1a-Cre) line, wherein Cre recombinase is expressed from the time of birth (Gerfen et al., 2013; Hoerder-Suabedissen et al., 2018). To generate these mice, Tg(Drd1a-cre)FK164Gsat/Mmucd (*Drd1a-Cre*; MMRRC)) and C57BL6-Snap25tm3mcw (*Snap25fl/fl)* animals were crossed to the tdTomato reporter strain B6;129S6-Gt(ROSA)26Sortm14(CAG-tdTomato)Hze/J (*Ai14*). The confirmation of the loss of regulated synaptic vesicular release from the Cre+ neuronal population was described in our previous publications (Marques-Smith et al., 2016; Hoerder-Suabedissen et al., 2019; Messore et al., 2025).

For baseline and sleep deprivation recordings, adult male L6b silenced animals (n=9), and Cre-negative adult male littermate control animals (n=7) were used following conventional group sizes in similar studies (Yamagata et al., 2021). For the orexin infusion experiments, separate cohorts of n=8 adult male L6b silenced animals and n=5 adult male littermate control animals were used. For the circadian screen, separate cohorts of young adult male L6b silenced (n=6) and control animals (n=7) were used. Genotypes were confirmed by polymerase chain reaction (PCR) through Transnetyx (Transnetyx Inc, Cordova, Tennessee, USA).

Although the age at the start of sleep experiments was comparable between genotypes (controls 11±0.48 weeks vs L6b silenced 10.6±0.39 weeks t_(14)_=0.6966,p=0.4974), L6b silenced animals had a significantly lower body weight (controls 23±0.67 g vs L6b silenced 21±0.47 g, t_(14)_=2.530, p=0.024). A lower body weight compared to Cre negative control animals has also been previously observed in Rbp4-Cre;Snap25^fl/fl^;Ai14 ‘L5 silenced’ mice (Hoerder-Suabedissen et al., 2019; Krone et al., 2021).

### Circadian screen

The circadian screen in L6b silenced mice was performed as a part of initial phenotyping in a separate study. Animals were housed in individual cages located inside light-tight chambers to allow control of the light-dark schedule, with food and water *ad libitum*. Home cage activity was assessed using running wheels, wherein rotations were recorded with a 1-minute resolution in ClockLab (ActiMetrics Inc, Wilmette, Illinois, USA, version 6.1.02). Baseline rhythmicity was investigated with a 12h:12h light-dark schedule, with lights on at 5:00AM. After 21 days, the light schedule was changed to constant darkness for 10 days to assess the intrinsic active period duration (alpha). Next, animals were kept in constant light for 16 days to characterise properties of the circadian clock in constant conditions. Lastly, a 6-hour phase-advance protocol was applied, in which lights were turned off at 11:00AM, corresponding to 6h after light onset, then on at 11:00PM. Recordings continued for 10 days.

### Surgical procedures and postoperative care

Surgical implantation of EEG/EMG recording electrodes was carried out as previously described (Krone et al., 2021; McKillop et al., 2021). Bregma was localized and the positions of the electrodes were marked, with the frontal EEG 2 mm anteroposterior (AP), +2 mm to the right of the midline (ML), occipital EEG -3.5mm AP and +2.5mm ML. Holes were created with a microdrill at the marked positions and above the cerebellum for the reference electrode. Next, the EEG/EMG headstage was positioned with the bone screws into the holes. The headstage was secured with dental cement (Super bond, Prestige Dental Products Ltd, Bradford, UK). EMGs were implanted into the nuchal muscles and secured to the headstage with acrylic dental cement (Simplex Rapid, Associated Dental Products Ltd, Swindon, UK). The headstage weight was <10% of presurgical body weight in all animals. Postoperative analgesia was provided through jellies mixed with oral Metacam (meloxicam, 5 mg/kg, Boehringer Ingelheim Ltd., Bracknell, UK) as needed.

For the orexin infusion experiment, surgeries were extended with the unilateral implantation of a cannula in the right lateral cerebral ventricle. To insert the cannula, an additional hole was drilled (AP -0.3mm, ML +0.9mm, DV -2.3mm; coordinates based on (Suzuki et al., 2005)) in the skull, and the guide cannula (C315G/Spc, Plastics1, Protech International Inc., Boerne, Texas, USA) was inserted into the holder in the stereotactic frame and slowly positioned atop the dura. The cannula was then lowered to 2.3 mm below the dura. Silicone elastomer (Kwik-Sil, World Precision Instruments Inc., Sarasota, FL, USA) was applied to close the craniotomy. The guide cannula was secured with dental cement (Super bond, Prestige Dental Products Ltd, Bradford, UK), and a dummy cannula (C315DC/Spc, Plastics1) was inserted into the guide cannula to prevent infection and bore occlusion. Dummies were manually moved in the cannulas every few days to habituate animals to the procedure and to ensure clean cannula bores.

### Recording setup

After recovery from surgery, animals were transferred to the recording room. Animals were housed in individual custom-made Plexiglas cages (20x32x35cm^3^) with bedding and nesting material under a controlled light-dark cycle (9AM light on, 9PM lights off). Food was supplied *ad libitum* on the floor of the cage and water was provided in a bottle. Two Plexiglas cages were placed in each sound-attenuated recording chamber (Campden Instruments, Loughborough, UK). Recording chambers were lit with an LED strip attached to the top of the chamber (light levels 120-180 lux), which was connected to a timer on the same light-dark schedule as the room. Chambers were ventilated through a fan mounted on the side wall of the chamber which was electrically shielded with aluminium foil. A hole in the top of the chamber allowed for the insertion of EEG/EMG cables and positioning of webcams (Logitech, C270, Lausanne, Switzerland) for remote monitoring. Room temperature and humidity were maintained at 20 ± 1°C and 60 ± 10%, respectively.

### Preparation of orexin solutions

ORXA (Orexin A (human, mouse, rat) Trifluoroacetate salt, 4028262.0500, lot no 1068668, Bachem AG, Switzerland) was dissolved in sterile saline to a concentration of 0.3 nmol/μl or 0.15 nmol/μl, and ORXB (Orexin B (human) Trifluoroacetate salt, 4028263.0500, lot no 1000056801, Bachem AG) to a concentration of 0.3 nmol/μl under sterile conditions in a flow cabinet. Doses were determined based on previous studies involving orexin infusion in mice (Mobarakeh et al., 2005; Suzuki et al., 2005; Mieda et al., 2011) and the reported dissolvability of orexin according to product data sheets.

### Cannula system

Each mouse had a separate tubing system consisting of an internal needle (C315I/SPC, Plastics1) inserted into the implanted guide canula, which was connected to a Hamilton syringe (SYR 5 μl 75N, Hamilton company, Nevada, USA) via transparent tubing (C313CT, C313C, Plastics1). The Hamilton syringe was placed in a microinjector pump (Pump 11, Elite, Harvard Apparatus, Massachusetts, USA). Tubing was front-filled with 3-4 μl drug solution and back-filled with saline, and marks were made to allow monitoring of fluid movement. Infusion success was therefore verified by the movement of the Hamilton plunger and the fluid level of the drug solution as compared to markings made preceding the infusion.

### Experimental timeline baseline and sleep deprivation recording

After a day of habituation, animals were connected to EEG/EMG cables and signals were checked using the Synapse recording software setup (Synapse, Tucker Davis Technologies, Florida, USA). After a few days, a baseline recording was undertaken. Typically, experiments lasted 1-3 weeks with 2-3 days in between experimental days other than those which involved baseline recordings.

For sleep deprivation (SD), animals were kept awake from light onset for 6 hours, which is when mice in laboratory conditions are predominantly asleep. At light onset (9AM), the chambers were opened, and nests were removed from the cages. Animals were continuously observed by an experimenter for the full 6 hours, and when an animal seemed to be falling asleep based on posture and/or online EEG/EMG traces, a novel object was introduced into the cage. This is a common SD procedure in rodents which keeps animals awake in an ethologically relevant and less-stressful manner than other SD methods (McKillop et al., 2018). After 6 hours, objects were taken out of the cages, nests were reintroduced, and chambers were closed.

### Experimental procedure orexin infusion

In the orexin infusion experiment, animals were moved into the Plexiglas cages in the recording room 5-7 days after recovering from surgery. Animals were connected to EEG/EMG cables a day later, then allowed to habituate to the new environment for several days before experiments commenced. The first day involved a baseline recording, wherein animals were only briefly disturbed for inspection and handling at 9AM. On each infusions day, chambers were opened at 9AM and animals were connected to the cannula system. Next, 2 μl of orexin or vehicle were administered at a rate of 1 μl/min within 55 min of light onset. After the infusion was complete, the system was left in place for an additional 5 minutes to allow the remaining solution to diffuse from the internal needle, and then the dummy cannulas were re-inserted, and mice were left undisturbed. The higher dose of ORXA was expected to give the strongest effect, so this infusion and a vehicle infusion were prioritised for the first two infusions in counterbalanced order. Additional infusions were administered longitudinally, with ORXB being counterbalanced with the lower dose of ORXA for the third and fourth infusion. There were 2-4 non-infusion days between consecutive infusion days. Infusion order was swapped for one individual animal for technical reasons.

### Data acquisition

EEG/EMG head stages were custom-made from 8-pin 90 degrees connectors (Pinnacle Technology Inc, Kansas, USA, model 8415-SM) to which stainless steel wires were attached that were wrapped around a stainless-steel bone screw (Fine Science Tools, 19010-10, InterFocus Ltd, Cambridge, UK) for the EEG electrodes; for the EMG electrodes, stainless steel wires were folded into a loop that was soldered into a smooth blob, and then attached to the 8-pin connector. Connections were conductivity-tested and electrically isolated with a layer of dental cement (Simplex Rapid dental acrylic cement (Associated Dental Products Ltd, Swindon, UK).

Data was acquired and stored locally using a 128-channel neurophysiology recording system with an RZ2 processor (Tucker Davis Technologies (TDT), Florida, USA) connected to a computer installed with TDT Synapse software (Tucker Davis Technologies). Custom-made EEG/EMG cables connected the headstage to a splitter-box in a configuration for analogue referencing (frontal-cerebellar, occipital-cerebellar, a backup derivation frontal-occipital, and EMG1-EMG2). EEG/EMG signals were preamplified by a PZ5 Neurodigitizer system (TDT), sampled at 1017.3 Hz (for baseline/sleep deprivation experiment recordings) or 305 Hz (orexin infusion experiment recordings), and processed with an anti-aliasing filter at 45% of the sampling rate.

### Signal processing

Blinded raw data from the circadian screen was analysed in Clocklab, then further processed in MATLAB (v2022a, The MathWorks Inc., Massachusetts, USA) and GraphPad Prism (v.9.3.1, GraphPad Software, Inc., California, USA). EEG and EMG signals were processed offline using custom-written MATLAB scripts. This involved first resampling signals to 256 Hz, then filtering to 0.3 Hz-100 Hz for EEG signals and 3-100 Hz for EMG signals using a Chebyshev type II filter. Resampled and filtered signals were then converted into European Data Format (.edf) files and blinded for manual assignment of vigilance states in 4-seconds epochs in SleepSign (Kissei Comptech Co Ltd, Nagano, Japan). Vigilance states were scored as non-rapid eye movement (NREM) sleep, rapid eye movement (REM) sleep, or Wakefulness (W) based on visual inspection of frontal and occipital EEG and EMG traces. Brief episodes of movements during NREM sleep (≤ 16 seconds) and REM sleep (≤ 8 seconds) were scored as movement rather than wakefulness and not considered a termination of sleep episodes. Minimum NREM sleep and Wake episode duration was defined as 1 minute and minimum REM sleep episode duration was defined as 16 seconds. Epochs with large movement and other noise were scored as artefacts of the respective vigilance state, which were included in vigilance state analysis but not spectral analysis. For spectral analysis, Fourier transforms were calculated from raw data using the pwelch function in MATLAB.

### Excluded data

i) Circadian screen. During the 12h:12h light-dark schedule, the first two days were excluded for all animals due to insufficient entrainment, and the last day was excluded for all animals because the data was incomplete. For both the constant dark and the constant light paradigm, one day was excluded for all animals due to missing data. For the constant light condition, data from one control animal was excluded for technical reasons. ii) Baseline and sleep deprivation recording. EEG and EMG channels of low quality (excessive noise and artifacts) were excluded from analysis (the occipital EEG channel in one control animal for both the baseline and SD day, and the occipital EEG channel of one additional control animal and a L6b silenced animal were excluded for the SD day only). In two (different) animals, EMG signals were of insufficient quality to reliably score bouts of movement, so these animals were excluded from analysis of brief awakenings. Epochs involving large movements and other noise were scored as artefacts of the respective vigilance state, which were included in vigilance state analysis but not EEG power spectral analysis. The percentage of artefact epochs across the 24-hour file was comparable between genotypes (controls = 2.80±1.43%, L6b silenced = 2.09±0.729%, t(14)=0.4706, p=0.65). iii) Orexin infusion experiment. We decided not to include a comparison of wake EEG spectra after orexin infusion because of the high variability in the duration of induced waking and the overall limited data set, primarily resulting from the exclusion of several animals for technical reasons affecting signal quality. Visual inspection of the remaining data of sufficient quality did not reveal obvious differences between genotypes; however, the sample size was insufficient to draw strong conclusions. Future studies will therefore be required to directly compare the effects of orexin on wake EEG activity between control and layer 6b-silenced mice.

### Statistics

Statistical analyses were conducted using GraphPad Prism. Statistical significance was defined as p ≤ α = 0.05. Interlayer density analyses employed two-way ANOVA, with post-hoc pairwise tests corrected for multiple comparisons using the Benjamini, Krieger and Yekutieli method. For EEG/EMG and sleep-wake state analyses, differences between L6b silenced and control animals across all metrics involving a single value per animal were evaluated using unpaired two-tailed t-tests, except for NREM sleep latency which was tested with a Mann Whitney U test. When there were multiple values per animal, genotype groups were compared using two-way Analyses of Variance (ANOVA) or with a Mixed Effects ANOVA when there were missing values. The Geisser-Greenhouse method was applied to correct for non-sphericity (F-test). Significant two-way interactions were followed up with post-hoc comparisons, applying the Bonferroni correction (0.05/number of observations) in all instances except for post-hoc comparisons of EEG spectra (no correction was applied, as adjusting for 119 frequency bins would be too conservative). Post hoc differences in frequency bins are only reported if two or more consecutive bins showed a difference (a range of >0.5 Hz). For EEG spectral analysis, data from individual animals were either log-transformed or normalised to baseline spectra before statistical testing. To investigate the acute effects of orexin, sleep architecture following orexin infusion was analysed starting at the time of each infusion and ending 3 hours thereafter (2700 epochs of 4 seconds each). Two doses of ORXA (0.3 nmol and 0.6 nmol) were administered, but as the effects of the lower dose appeared to be consistent with the effects of the higher dose, subsequent descriptions for some analyses focus only on the higher dose of ORXA.

### Histology

After experiment completion, animals were deeply anaesthetised with a lethal dose of pentobarbitone and perfused transcardially with 0.1 M Phosphate Buffered Saline (PBS), followed by 4% formaldehyde (F8775; Sigma-Aldrich) in 0.1 M PBS. Head stages (including cannulas) were removed, then brains were extracted for post-fixation in 4% PFA for 24 hours at 4 °C before being transferred to 0.05% PBS-azide (PBSA) for long-term storage at 4 °C. Brains were sliced into 50μm-thick coronal sections using a vibroslicer (Leica VT1000S). Free floating sections were collected and stored in PBSA. For verification of cannula locations, nuclei were stained with 1:1000 4′,6- diamidino-2-phenylindole (DAPI, Invitrogen, D1306) for visualisation of tissue structure. For determination of Drd1a cell density (Fig. 1), sections were incubated with primary antibodies (rabbit α-Cplx3 to distinguish between L6a and L6b (1:1000, Synaptic Systems #122302), mouse α-Tbr1 to identify the L5-6 boundary (1:500, ProteinTech #66564-1-LG) in 5% donkey serum, 0.3% Triton-X, 0.1M PBS) at 4°C for 48hrs. After 3x10 min washes, sections were incubated with secondary antibodies (donkey α-Rb (1:500, Invitrogen #A21206, 488nm), donkey α-mouse (1:500, Invitrogen #A32787, 647nm) in 5% donkey serum, 0.3% Triton-X, 0.1M PBS) for 2 hrs at room temperature. After another set of 3×10min washes, sections were incubated with 1:1000 4′,6-diamidino-2- phenylindole (DAPI) for 30mins at room temperature and washed 2×10min in 0.1M PBS. Sections were mounted using mounting medium (Fluorsave, VWR, Lutterworth UK) on microscope slides (Thermoscientific superfrost, RF12312108, Loughborough, UK) with cover slips prior to imaging (VWR coverglasses, 24 x 50mm, 531-0146, Lutterworth UK).

### Imaging

Images for cell quantifications were obtained using 20X spinning-disk confocal microscopy (Olympus SpinSR SoRa) and pre-processed by maximum-intensity z-stack projection, auto-brightness thresholding, and background subtraction using a custom Fiji/ImageJ .ijm script. Outputs were imported into QuPath (v0.4.2) (Bankhead et al., 2017) for cell segmentation. Resulting detections and downsampled images were additionally processed using another custom Fiji/ImageJ.ijm script (i.e., binary masks application, pixel inversion, and file type conversion to RGB .pngs), merged for atlas registration (2017 CCFv3) (Wang et al., 2020), and analysed using the QUINT pipeline (Yates et al., 2019). Software versions employed as a part of the QUINT pipeline include QuickNII (RRID:SCR_016854) (Puchades et al., 2019), VisuAlign (v0.8 RRID:SCR_017978), and Nutil (generating object reports with minimum pixel size = 4, point cloud density = 4, without object splitting, and extracting all coordinates) (Groeneboom et al., 2020). A custom MATLAB script (R2022b) re-formatted Nutil outputs for statistical analysis and graphical representation using GraphPad Prism. For the orexin experiment, brain sections were imaged with an epifluorescence microscope mounted with a camera (Leica Digital Module R, (DMR) with DFC500) or laser scanning confocal microscope (Zeiss LSM710) to confirm cannulas successfully penetrated the lateral ventricle. Images were processed in Fiji/ImageJ (FIJI, Schindelin et al 2012, 2.9.0 v1.54b) using background subtraction, brightness, and contrast adjustments, and merging of images from different fluorescent channels. Further processing, such as adding labels and indicating histological or anatomical boundaries, were carried out in Inkscape (Inkscape Project. (2020). *Inkscape*. Retrieved from https://inkscape.org).

## Acknowledgements

St John’s College Research Centre Grant on L6b to EM and ZM

ZM and Ed Mann MRC Grant MR/W029073/1; VV and ZM BBSRC grant BB/X008711/1

Medical Research Council (UK) grant MR/S01134X/1 (VV)

VV is supported by The MRC grant MR/V013238/1 and The Wellcome Trust grant 227093/Z/23/Z

EM was funded by the O’Sullivan Family Graduate Scholarship

LG was funded by a WT studentship.

MM was funded by the Rhodes Scholarship, Clarendon Scholarship, and Goodger and Schorstein Scholarship (University of Oxford Department of Physiology Anatomy and Genetics).

SW was funded by an NC3Rs studentship (NC/S001689/1)

AC was funded by a BBSRC studentship (BB/M011224/1)

HA was funded by the Wellcome trust through a postdoctoral fellowship (206500/Z/17/Z).

LK was supported by the Wellcome Trust through a doctoral studentship (203971/Z/16/Z) and postdoctoral fellowship (224083/Z/21/Z) as well as the Staines Medical Research Fellowship at Exeter College Oxford.

## Data availability

https://doi.org/10.6084/m9.figshare.31375441

## Notes

### Competing Interest Statement

The authors have declared no competing interest.

### Summary of Updates

The Drd1a Cre mouse model used (FK164) has a relatively selective expression of Drd1a Cre in cortex, but indeed some expression is seen subcortically. This is an acknowledged limitation which is now explicitly addressed in the revised manuscript. In our previous publications, we showed confirmation of the loss of regulated synaptic vesicle release from the Cre-positive neuronal population (Marques-Smith et al., 2016; Hoerder-Suabedissen et al., 2018; Messore et al., 2024). This has now been described in the revised manuscript. We have now carefully revised the spectral results and implemented a consistent approach in statistical reporting and spectral plots. We have updated Supplementary Table T1, Figure 3a and S2 to ensure that all statistics are presented consistently throughout the manuscript, i.e. with two-way ANOVAs and accompanying posthoc tests. We have adjusted markers to reflect the results from posthoc tests after two-way ANOVAs. There were 2-4 non-infusions days between infusions. We have added this information to methods. We have adjusted the wording in the main text to reflect more precisely which comparisons are shown in the figures. We now acknowledge the limitations that the ICV route of orexin administration cannot guarantee that only cortical Drd1a-Cre expressing neurons are reached by orexin, and the Drd1a-Cre driver line is highly selective but not entirely specific for layer 6b neurons We now provide the rationale for using only male mice in the methods section.

