## Supplementary Figures and Tables 1-3 for "Cortical layer 6b mediates state-dependent changes in brain activity and effects of orexin on waking and sleep"

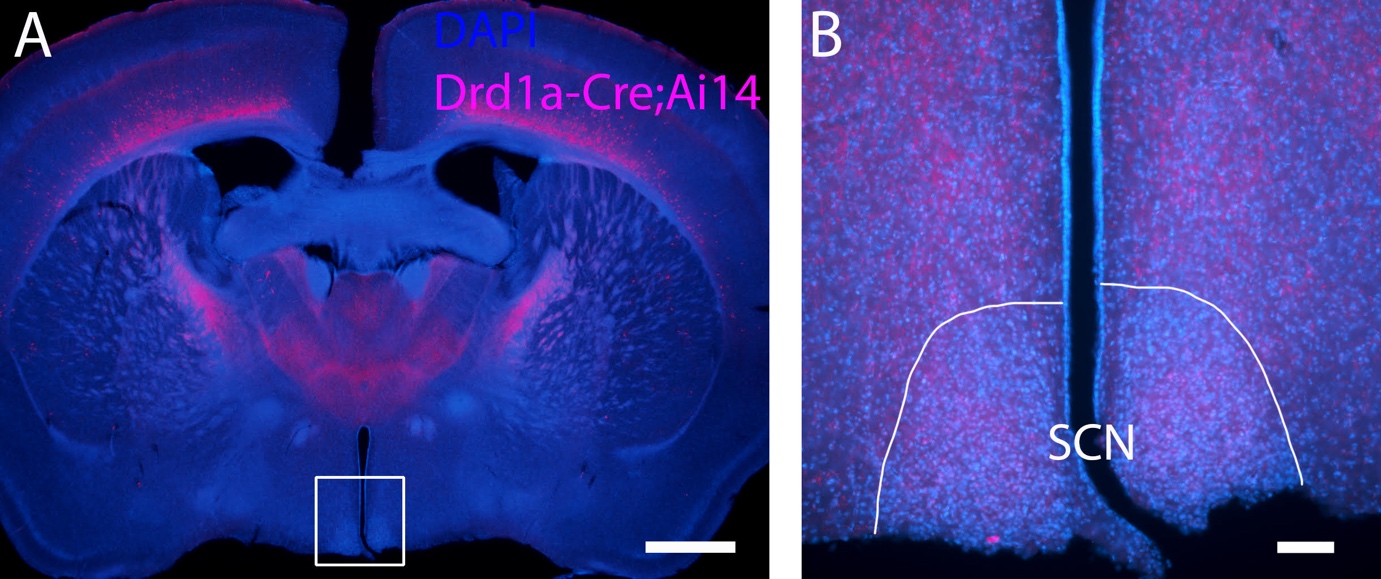

**Figure S1. Absence of Drd1a-Cre positive cells in the suprachiasmatic nucleus**

Representative images from an individual Drd1a-Cre; Ai14 animal. Tissue structure is demarcated with DAPI (blue) and Drd1a-Cre cells are marked by TdTom expression (magenta).

a. Overview (1.6X) with white square indicating the area magnified in b. Scale bar, 1000 µm.

b. No Cre expressing cells are present in the suprachiasmatic nucleus (SCN). Scale bar, 100 µm.

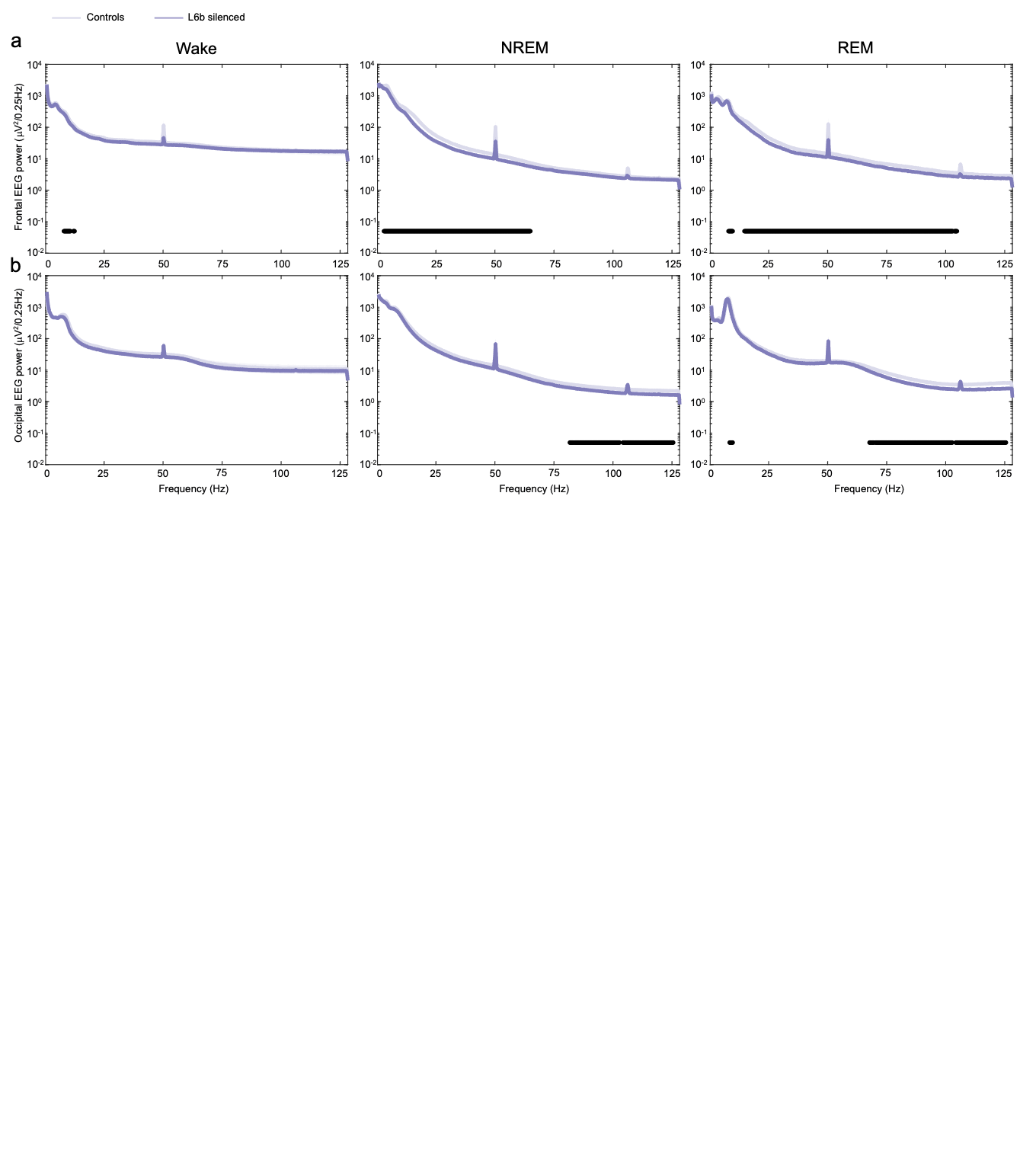

**Figure S2. Spectral power density across wake, NREM and REM across the full frequency range**

a. Frontal EEG power during the respective vigilance states in L6b silenced (n=9) and control (n=7) animals. Filled circles depict significantly differences between genotypes in 0.25-Hz bin power spectra in posthoc tests when two-way ANOVAs showed a significant genotype x frequency interaction.
b. Occipital EEG during the respective vigilance states in L6b silenced (n=9) and control (n=6) animals. Filled circles depict significantly differences between genotypes in 0.25-Hz bin power spectra in posthoc tests when two-way ANOVAs showed a significant genotype x frequency interaction.

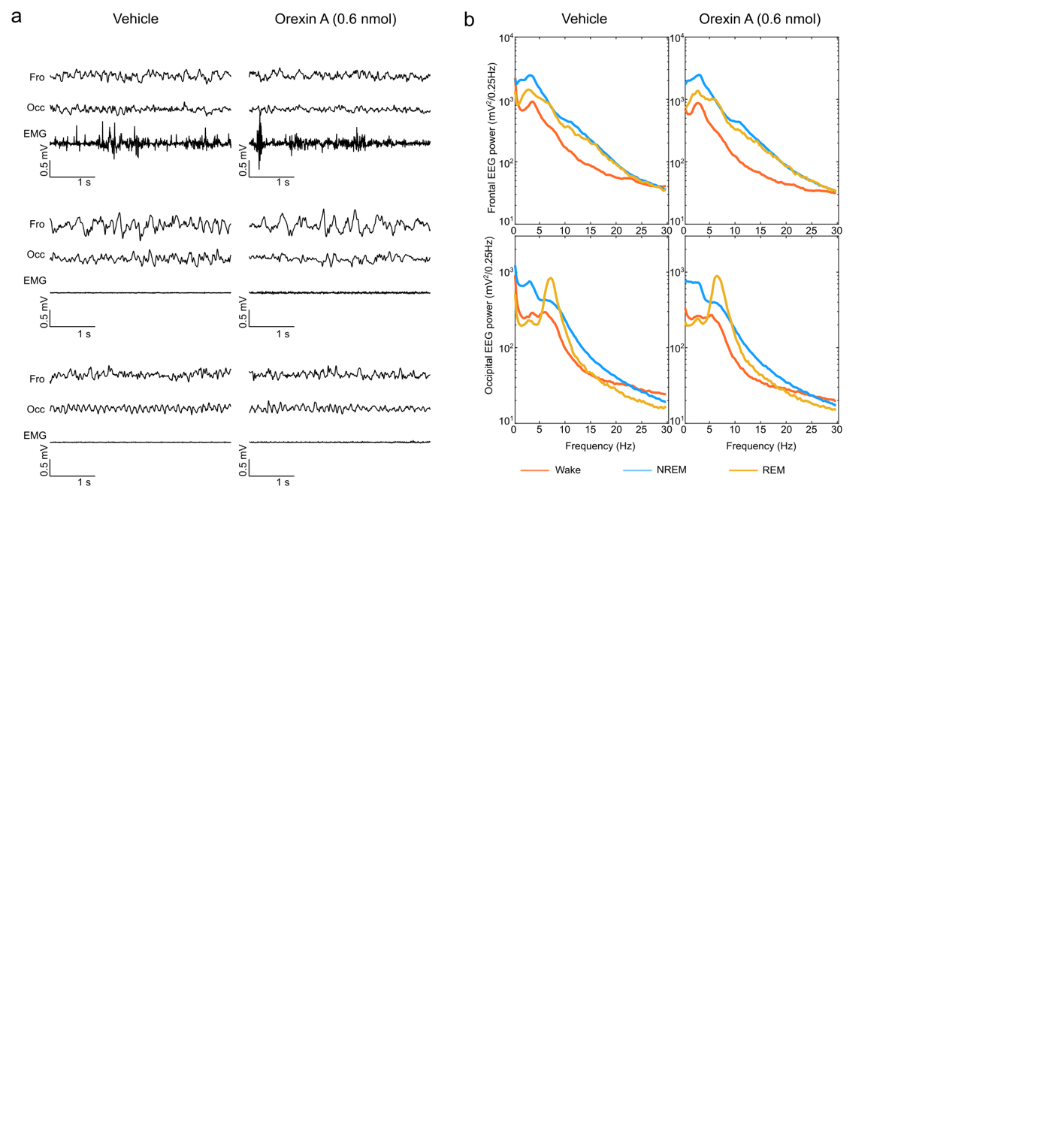

**Figure S3. Effects of orexin A on EEG spectral power in a representative individual animal**

a. Representative epoch of the frontal EEG (top), occipital EEG (middle) and EMG (bottom) channel in a representative control animal during wake, NREM and REM. The left column shows traces after vehicle infusion, the right column shows traces after the higher dose (0.6 nmol) of ORXA infusion.

b. EEG power spectra for the frontal EEG (top) and occipital EEG (bottom) after vehicle infusion (left column) and ORXA (0.6 nmol) infusion in a representative control animal, averaged across all wake, NREM and REM epochs during 24 hours.

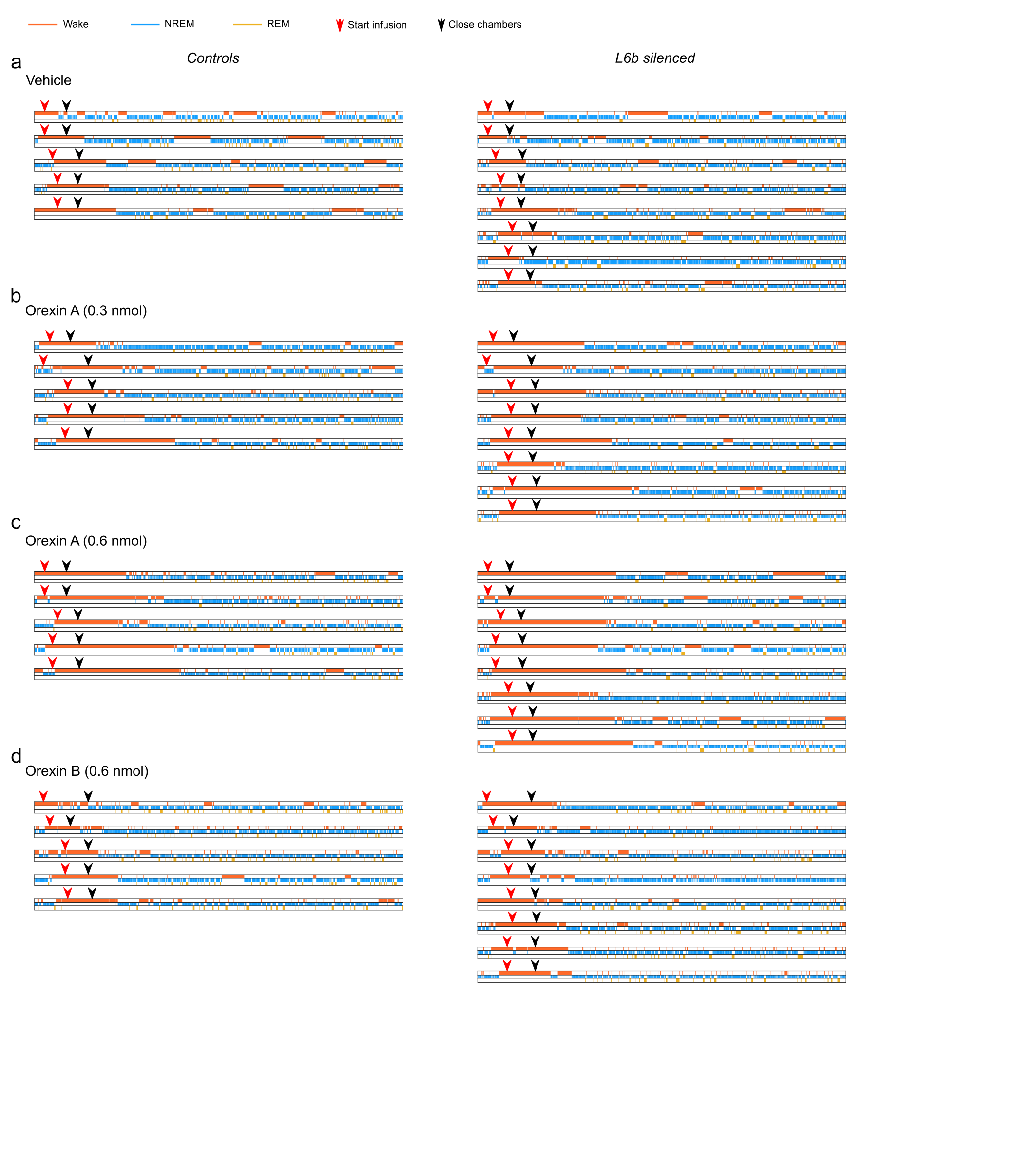
**Figure S4. Hypnograms off all animals after the different infusions**

Each row represents a 6-hour hypnogram from light onset for an individual animal after (a) vehicle infusion, (b) infusion of the lower dose of orexin A (0.3 nmol), (c) infusion of the higher dose of orexin A (0.6 nmol) and (d) infusion of orexin B (0.6 nmol). The red arrowheads indicate the start of the infusion, the black arrowheads represent the closure of the recording chamber. Hypnograms are split out per genotype, with controls on the left and L6b silenced animals on the right.

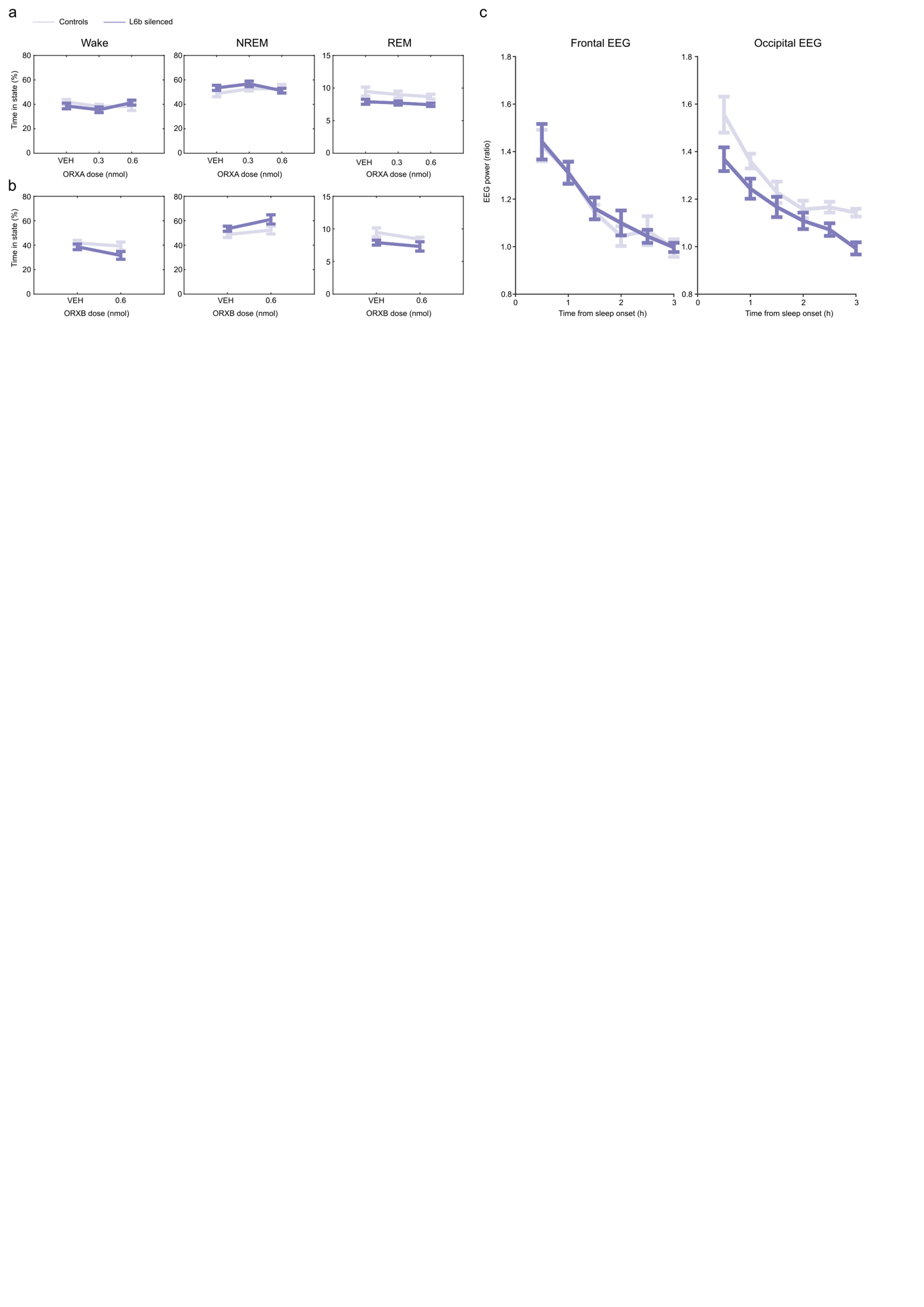
**Figure S5 Homeostatic regulation of sleep after orexin infusion**

a. Time spent in wake, NREM and REM after vehicle (saline) and ORXA infusion (dose 0.3 nmol and 0.6 nmol respectively).

b. Time spent in wake, NREM and REM after vehicle (saline) and ORXB (dose 0.6 nmol) infusion.

c. Slow wave activity rebound during NREM sleep following ORXA infusion (0.6 nmol) in the frontal EEG and occipital EEG. Controls n=5, L6b silenced n=8.

**Supplementary Table T1. Inter-layer pairwise comparisons**

*Method = Two-stage linear step-up procedure of Benjamini Krieger and Yekutieli; number of families = 4; number of comparisons per family = 21; significance defined at q/p < α = 0.05.*

| **Cortical layer**  **comparison:**  **Main results** | **M1** | | | **S1** | | |
| --- | --- | --- | --- | --- | --- | --- |
|  | **Predicted mean difference (LS)** | **q value** | **p value** | **Predicted mean difference (LS)** | **q value** | **p value** |
| L6b vs. L6a | 0.00065 | <0.0001 | <0.0001 | 0.000665 | <0.0001 | <0.0001 |
| L6b vs. L5 | 0.001448 | <0.0001 | <0.0001 | 0.000949 | <0.0001 | <0.0001 |
| L6b vs. L4 | 0.001461 | <0.0001 | <0.0001 | 0.000951 | <0.0001 | <0.0001 |
| L6b vs. L3 | 0.001456 | <0.0001 | <0.0001 | 0.000952 | <0.0001 | <0.0001 |
| L6b vs. L2 | 0.001456 | <0.0001 | <0.0001 | 0.000952 | <0.0001 | <0.0001 |
| L6b vs. L1 | 0.001461 | <0.0001 | <0.0001 | 0.000953 | <0.0001 | <0.0001 |
| L6a vs. L5 | 0.000798 | <0.0001 | <0.0001 | 0.000284 | <0.0001 | <0.0001 |
| L6a vs. L4 | 0.000811 | <0.0001 | <0.0001 | 0.000286 | <0.0001 | <0.0001 |
| L6a vs. L3 | 0.000806 | <0.0001 | <0.0001 | 0.000287 | <0.0001 | <0.0001 |
| L6a vs. L2 | 0.000806 | <0.0001 | <0.0001 | 0.000287 | <0.0001 | <0.0001 |
| L6a vs. L1 | 0.000811 | <0.0001 | <0.0001 | 0.000288 | <0.0001 | <0.0001 |
| L5 vs. L4 | 1.35E-05 | 0.5 | 0.8105 | 2.33E-06 | 0.5 | 0.967 |
| L5 vs. L3 | 8.59E-06 | 0.5 | 0.8787 | 3.28E-06 | 0.5 | 0.9535 |
| L5 vs. L2 | 8.59E-06 | 0.5 | 0.8787 | 3.28E-06 | 0.5 | 0.9535 |
| L5 vs. L1 | 1.35E-05 | 0.5 | 0.8105 | 4.08E-06 | 0.5 | 0.9423 |
| L4 vs. L3 | -4.91E-06 | 0.5 | 0.9305 | 9.50E-07 | 0.5 | 0.9865 |
| L4 vs. L2 | -4.91E-06 | 0.5 | 0.9305 | 9.50E-07 | 0.5 | 0.9865 |
| L4 vs. L1 | 0 | 0.5 | >0.9999 | 1.75E-06 | 0.5 | 0.9753 |
| L3 vs. L2 | 1.08E-19 | 0.5 | >0.9999 | 0 | 0.5 | >0.9999 |
| L3 vs. L1 | 4.91E-06 | 0.5 | 0.9305 | 7.95E-07 | 0.5 | 0.9887 |
| L2 vs. L1 | 4.91E-06 | 0.5 | 0.9305 | 7.95E-07 | 0.5 | 0.9887 |
| **Cortical layer**  **comparison:**  **Main results** | **V1** | | | **PFC** | | |
|  | **Predicted mean difference (LS)** | **q value** | **p value** | **Predicted mean difference (LS)** | **q value** | **p value** |
| L6b vs. L6a | 0.000724 | <0.0001 | <0.0001 | 0.000496 | <0.0001 | <0.0001 |
| L6b vs. L5 | 0.000898 | <0.0001 | <0.0001 | 0.000915 | <0.0001 | <0.0001 |
| L6b vs. L4 | 0.000899 | <0.0001 | <0.0001 | 0.000923 | <0.0001 | <0.0001 |
| L6b vs. L3 | 0.000897 | <0.0001 | <0.0001 | 0.000922 | <0.0001 | <0.0001 |
| L6b vs. L2 | 0.000897 | <0.0001 | <0.0001 | 0.000922 | <0.0001 | <0.0001 |
| L6b vs. L1 | 0.000899 | <0.0001 | <0.0001 | 0.000923 | <0.0001 | <0.0001 |
| L6a vs. L5 | 0.000174 | 0.0454 | 0.0304 | 0.000419 | <0.0001 | <0.0001 |
| L6a vs. L4 | 0.000175 | 0.0454 | 0.0296 | 0.000427 | <0.0001 | <0.0001 |
| L6a vs. L3 | 0.000173 | 0.0454 | 0.0317 | 0.000426 | <0.0001 | <0.0001 |
| L6a vs. L2 | 0.000173 | 0.0454 | 0.0317 | 0.000426 | <0.0001 | <0.0001 |
| L6a vs. L1 | 0.000175 | 0.0454 | 0.0296 | 0.000427 | <0.0001 | <0.0001 |
| L5 vs. L4 | 8.37E-07 | 0.75 | 0.9916 | 7.98E-06 | 0.5 | 0.897 |
| L5 vs. L3 | -1.43E-06 | 0.75 | 0.9856 | 6.65E-06 | 0.5 | 0.9141 |
| L5 vs. L2 | -1.43E-06 | 0.75 | 0.9856 | 6.65E-06 | 0.5 | 0.9141 |
| L5 vs. L1 | 8.37E-07 | 0.75 | 0.9916 | 7.44E-06 | 0.5 | 0.904 |
| L4 vs. L3 | -2.27E-06 | 0.75 | 0.9772 | -1.33E-06 | 0.5 | 0.9828 |
| L4 vs. L2 | -2.27E-06 | 0.75 | 0.9772 | -1.33E-06 | 0.5 | 0.9828 |
| L4 vs. L1 | 0 | 0.75 | >0.9999 | -5.41E-07 | 0.5 | 0.993 |
| L3 vs. L2 | -1.08E-19 | 0.75 | >0.9999 | -5.42E-20 | 0.5 | >0.9999 |
| L3 vs. L1 | 2.27E-06 | 0.75 | 0.9772 | 7.89E-07 | 0.5 | 0.9898 |
| L2 vs. L1 | 2.27E-06 | 0.75 | 0.9772 | 7.89E-07 | 0.5 | 0.9898 |

*Additional test details are as follows: For M1 and S1, N1 = N2 = 6 and SE of differences = 0.00005616 with DF = 112; for V1, N1 = N2 = 3 and SE of differences = 0.00007942 with DF = 112; and for PFC, N1 = N2 = 5 and SE of differences = 0.00006152 with DF = 112.*

| **Cortical layer**  **comparison:**  **Test details** | **M1** | | | **S1** | | |
| --- | --- | --- | --- | --- | --- | --- |
|  | **Predicted mean 1 (LS)** | **Predicted mean 2 (LS)** | **t value** | **Predicted mean 1 (LS)** | **Predicted mean 2 (LS)** | **t value** |
| L6b vs. L6a | 0.001461 | 0.000811 | 11.57 | 0.000954 | 0.000289 | 11.84 |
| L6b vs. L5 | 0.001461 | 1.35E-05 | 25.78 | 0.000954 | 5.00E-06 | 16.9 |
| L6b vs. L4 | 0.001461 | 0 | 26.02 | 0.000954 | 2.67E-06 | 16.94 |
| L6b vs. L3 | 0.001461 | 4.91E-06 | 25.93 | 0.000954 | 1.72E-06 | 16.96 |
| L6b vs. L2 | 0.001461 | 4.91E-06 | 25.93 | 0.000954 | 1.72E-06 | 16.96 |
| L6b vs. L1 | 0.001461 | 0 | 26.02 | 0.000954 | 9.25E-07 | 16.97 |
| L6a vs. L5 | 0.000811 | 1.35E-05 | 14.20 | 0.000289 | 5.00E-06 | 5.057 |
| L6a vs. L4 | 0.000811 | 0 | 14.44 | 0.000289 | 2.67E-06 | 5.099 |
| L6a vs. L3 | 0.000811 | 4.91E-06 | 14.35 | 0.000289 | 1.72E-06 | 5.116 |
| L6a vs. L2 | 0.000811 | 4.91E-06 | 14.35 | 0.000289 | 1.72E-06 | 5.116 |
| L6a vs. L1 | 0.000811 | 0 | 14.44 | 0.000289 | 9.25E-07 | 5.13 |
| L5 vs. L4 | 1.35E-05 | 0 | 0.24 | 5.00E-06 | 2.67E-06 | 0.04149 |
| L5 vs. L3 | 1.35E-05 | 4.91E-06 | 0.15 | 5.00E-06 | 1.72E-06 | 0.05841 |
| L5 vs. L2 | 1.35E-05 | 4.91E-06 | 0.15 | 5.00E-06 | 1.72E-06 | 0.05841 |
| L5 vs. L1 | 1.35E-05 | 0 | 0.24 | 5.00E-06 | 9.25E-07 | 0.07256 |
| L4 vs. L3 | 0 | 4.91E-06 | 0.09 | 2.67E-06 | 1.72E-06 | 0.01692 |
| L4 vs. L2 | 0 | 4.91E-06 | 0.09 | 2.67E-06 | 1.72E-06 | 0.01692 |
| L4 vs. L1 | 0 | 0 | 0.00 | 2.67E-06 | 9.25E-07 | 0.03107 |
| L3 vs. L2 | 4.91E-06 | 4.91E-06 | 0.00 | 1.72E-06 | 1.72E-06 | 0 |
| L3 vs. L1 | 4.91E-06 | 0 | 0.09 | 1.72E-06 | 9.25E-07 | 0.01416 |
| L2 vs. L1 | 4.91E-06 | 0 | 0.09 | 1.72E-06 | 9.25E-07 | 0.01416 |
| **Cortical layer**  **comparison:**  **Test details** | **V1** | | | **PFC** | | |
|  | **Predicted mean 1 (LS)** | **Predicted mean 2 (LS)** | **t value** | **Predicted mean 1 (LS)** | **Predicted mean 2 (LS)** | **t value** |
| L6b vs. L6a | 0.000899 | 0.000175 | 9.116 | 0.000923 | 0.000427 | 8.063 |
| L6b vs. L5 | 0.000899 | 8.37E-07 | 11.31 | 0.000923 | 7.98E-06 | 14.87 |
| L6b vs. L4 | 0.000899 | 0 | 11.32 | 0.000923 | 0 | 15 |
| L6b vs. L3 | 0.000899 | 2.27E-06 | 11.29 | 0.000923 | 1.33E-06 | 14.98 |
| L6b vs. L2 | 0.000899 | 2.27E-06 | 11.29 | 0.000923 | 1.33E-06 | 14.98 |
| L6b vs. L1 | 0.000899 | 0 | 11.32 | 0.000923 | 5.41E-07 | 14.99 |
| L6a vs. L5 | 0.000175 | 8.37E-07 | 2.193 | 0.000427 | 7.98E-06 | 6.811 |
| L6a vs. L4 | 0.000175 | 0 | 2.203 | 0.000427 | 0 | 6.941 |
| L6a vs. L3 | 0.000175 | 2.27E-06 | 2.175 | 0.000427 | 1.33E-06 | 6.919 |
| L6a vs. L2 | 0.000175 | 2.27E-06 | 2.175 | 0.000427 | 1.33E-06 | 6.919 |
| L6a vs. L1 | 0.000175 | 0 | 2.203 | 0.000427 | 5.41E-07 | 6.932 |
| L5 vs. L4 | 8.37E-07 | 0 | 0.01054 | 7.98E-06 | 0 | 0.1297 |
| L5 vs. L3 | 8.37E-07 | 2.27E-06 | 0.01804 | 7.98E-06 | 1.33E-06 | 0.1081 |
| L5 vs. L2 | 8.37E-07 | 2.27E-06 | 0.01804 | 7.98E-06 | 1.33E-06 | 0.1081 |
| L5 vs. L1 | 8.37E-07 | 0 | 0.01054 | 7.98E-06 | 5.41E-07 | 0.1209 |
| L4 vs. L3 | 0 | 2.27E-06 | 0.02858 | 0 | 1.33E-06 | 0.02162 |
| L4 vs. L2 | 0 | 2.27E-06 | 0.02858 | 0 | 1.33E-06 | 0.02162 |
| L4 vs. L1 | 0 | 0 | 0 | 0 | 5.41E-07 | 0.008794 |
| L3 vs. L2 | 2.27E-06 | 2.27E-06 | 1.4E-15 | 1.33E-06 | 1.33E-06 | 8.81E-16 |
| L3 vs. L1 | 2.27E-06 | 0 | 0.02858 | 1.33E-06 | 5.41E-07 | 0.01283 |
| L2 vs. L1 | 2.27E-06 | 0 | 0.02858 | 1.33E-06 | 5.41E-07 | 0.01283 |

**Supplementary Table T2. Inter-region pairwise comparisons**

*Method = Two-stage linear step-up procedure of Benjamini Krieger and Yekutieli; number of families = 7; number of comparisons per family = 6; significance defined at q/p < α = 0.05.*

| **Cortical region comparison: Main results** | | **Predicted mean difference (LS)** | **q value** | **p value** |
| --- | --- | --- | --- | --- |
| **L6b** | M1 vs. S1 | 0.000507 | <0.0001 | <0.0001 |
|  | M1 vs. V1 | 0.000562 | <0.0001 | <0.0001 |
|  | M1 vs. PFC | 0.000538 | <0.0001 | <0.0001 |
|  | S1 vs. V1 | 5.50E-05 | 0.3352 | 0.4256 |
|  | S1 vs. PFC | 3.10E-05 | 0.3778 | 0.5997 |
|  | V1 vs. PFC | -2.40E-05 | 0.3865 | 0.7361 |
| **L6a** | M1 vs. S1 | 0.000522 | <0.0001 | <0.0001 |
|  | M1 vs. V1 | 0.000636 | <0.0001 | <0.0001 |
|  | M1 vs. PFC | 0.000384 | <0.0001 | <0.0001 |
|  | S1 vs. V1 | 0.000114 | 0.0175 | 0.1002 |
|  | S1 vs. PFC | -0.00014 | 0.0044 | 0.0209 |
|  | V1 vs. PFC | -0.00025 | 0.0001 | 0.0006 |
| **L5** | M1 vs. S1 | 8.50E-06 | >0.9999 | 0.88 |
|  | M1 vs. V1 | 1.27E-05 | >0.9999 | 0.8543 |
|  | M1 vs. PFC | 5.52E-06 | >0.9999 | 0.9255 |
|  | S1 vs. V1 | 4.16E-06 | >0.9999 | 0.9518 |
|  | S1 vs. PFC | -2.98E-06 | >0.9999 | 0.9597 |
|  | V1 vs. PFC | -7.14E-06 | >0.9999 | 0.9201 |
| **L4** | M1 vs. S1 | -2.67E-06 | >0.9999 | 0.9622 |
|  | M1 vs. V1 | 0 | >0.9999 | >0.9999 |
|  | M1 vs. PFC | 0 | >0.9999 | >0.9999 |
|  | S1 vs. V1 | 2.67E-06 | >0.9999 | 0.9691 |
|  | S1 vs. PFC | 2.67E-06 | >0.9999 | 0.9639 |
|  | V1 vs. PFC | 0 | >0.9999 | >0.9999 |
| **L3** | M1 vs. S1 | 3.19E-06 | >0.9999 | 0.9548 |
|  | M1 vs. V1 | 2.64E-06 | >0.9999 | 0.9695 |
|  | M1 vs. PFC | 3.58E-06 | >0.9999 | 0.9516 |
|  | S1 vs. V1 | -5.50E-07 | >0.9999 | 0.9936 |
|  | S1 vs. PFC | 3.90E-07 | >0.9999 | 0.9947 |
|  | V1 vs. PFC | 9.40E-07 | >0.9999 | 0.9895 |
| **L2** | M1 vs. S1 | 3.19E-06 | >0.9999 | 0.9548 |
|  | M1 vs. V1 | 2.64E-06 | >0.9999 | 0.9695 |
|  | M1 vs. PFC | 3.58E-06 | >0.9999 | 0.9516 |
|  | S1 vs. V1 | -5.50E-07 | >0.9999 | 0.9936 |
|  | S1 vs. PFC | 3.90E-07 | >0.9999 | 0.9947 |
|  | V1 vs. PFC | 9.40E-07 | >0.9999 | 0.9895 |
| **L1** | M1 vs. S1 | -9.25E-07 | >0.9999 | 0.9869 |
|  | M1 vs. V1 | 0 | >0.9999 | >0.9999 |
|  | M1 vs. PFC | -5.41E-07 | >0.9999 | 0.9927 |
|  | S1 vs. V1 | 9.25E-07 | >0.9999 | 0.9893 |
|  | S1 vs. PFC | 3.84E-07 | >0.9999 | 0.9948 |
|  | V1 vs. PFC | -5.41E-07 | >0.9999 | 0.9939 |

*Additional test details are as follows: For SE of differences, M1 vs S1 = 5.62E-05, M1 vs V1 = S1 vs V1 = 6.88E-05, M1 vs PFC = S1 vs PFC = 5.89E-05, and V1 vs PFC = 7.10E-05; N = 6 for M1 and S1, N = 3 for V1, and N = 5 for PFC; and DF = 112 for all comparisons.*

| **Cortical region comparison: Test details** | | **Predicted mean 1 (LS)** | **Predicted mean 2 (LS)** | **t value** |
| --- | --- | --- | --- | --- |
| **L6b** | M1 vs. S1 | 0.001461 | 0.000954 | 9.028 |
|  | M1 vs. V1 | 0.001461 | 0.000899 | 8.171 |
|  | M1 vs. PFC | 0.001461 | 0.000923 | 9.134 |
|  | S1 vs. V1 | 0.000954 | 0.000899 | 0.7997 |
|  | S1 vs. PFC | 0.000954 | 0.000923 | 0.5263 |
|  | V1 vs. PFC | 0.000899 | 0.000923 | 0.3379 |
| **L6a** | M1 vs. S1 | 0.000811 | 0.000289 | 9.295 |
|  | M1 vs. V1 | 0.000811 | 0.000175 | 9.247 |
|  | M1 vs. PFC | 0.000811 | 0.000427 | 6.52 |
|  | S1 vs. V1 | 0.000289 | 0.000175 | 1.657 |
|  | S1 vs. PFC | 0.000289 | 0.000427 | 2.343 |
|  | V1 vs. PFC | 0.000175 | 0.000427 | 3.548 |
| **L5** | M1 vs. S1 | 1.35E-05 | 5.00E-06 | 0.1514 |
|  | M1 vs. V1 | 1.35E-05 | 8.37E-07 | 0.1841 |
|  | M1 vs. PFC | 1.35E-05 | 7.98E-06 | 0.09372 |
|  | S1 vs. V1 | 5.00E-06 | 8.37E-07 | 0.06053 |
|  | S1 vs. PFC | 5.00E-06 | 7.98E-06 | 0.05059 |
|  | V1 vs. PFC | 8.37E-07 | 7.98E-06 | 0.1006 |
| **L4** | M1 vs. S1 | 0 | 2.67E-06 | 0.04754 |
|  | M1 vs. V1 | 0 | 0 | 0 |
|  | M1 vs. PFC | 0 | 0 | 0 |
|  | S1 vs. V1 | 2.67E-06 | 0 | 0.03882 |
|  | S1 vs. PFC | 2.67E-06 | 0 | 0.04533 |
|  | V1 vs. PFC | 0 | 0 | 0 |
| **L3** | M1 vs. S1 | 4.91E-06 | 1.72E-06 | 0.0568 |
|  | M1 vs. V1 | 4.91E-06 | 2.27E-06 | 0.03838 |
|  | M1 vs. PFC | 4.91E-06 | 1.33E-06 | 0.06078 |
|  | S1 vs. V1 | 1.72E-06 | 2.27E-06 | 0.007997 |
|  | S1 vs. PFC | 1.72E-06 | 1.33E-06 | 0.006621 |
|  | V1 vs. PFC | 2.27E-06 | 1.33E-06 | 0.01323 |
| **L2** | M1 vs. S1 | 4.91E-06 | 1.72E-06 | 0.0568 |
|  | M1 vs. V1 | 4.91E-06 | 2.27E-06 | 0.03838 |
|  | M1 vs. PFC | 4.91E-06 | 1.33E-06 | 0.06078 |
|  | S1 vs. V1 | 1.72E-06 | 2.27E-06 | 0.007997 |
|  | S1 vs. PFC | 1.72E-06 | 1.33E-06 | 0.006621 |
|  | V1 vs. PFC | 2.27E-06 | 1.33E-06 | 0.01323 |
| **L1** | M1 vs. S1 | 0 | 9.25E-07 | 0.01647 |
|  | M1 vs. V1 | 0 | 0 | 0 |
|  | M1 vs. PFC | 0 | 5.41E-07 | 0.009185 |
|  | S1 vs. V1 | 9.25E-07 | 0 | 0.01345 |
|  | S1 vs. PFC | 9.25E-07 | 5.41E-07 | 0.00652 |
|  | V1 vs. PFC | 0 | 5.41E-07 | 0.007616 |

|  | **EEG** | **F(DFn, DFd)** | **p** | **Significant posthoc comparisons** |
| --- | --- | --- | --- | --- |
| **Wake** | Frontal | F(500,700)=1.235 | p=0.0004 | 8.0-10.75, 12-12.75 Hz |
|  | Occipital | F(500,6500)=0.6897 | p>0.9999 | 9.0-13.75, 14-15 Hz |
| **NREM** | Frontal | F(500,7000)=5.566 | p<0.0001 | 3.0-67.75 Hz |
|  | Occipital | F(500,6500)=1.131 | p=0.0274 | 84.25-128 Hz |
| **REM** | Frontal | F(500,7000)=1.563 | p<0.0001 | 8.0-10 Hz, 14.75-107.5 Hz |
|  | Occipital | F(500,6500)=6.284 | p<0.0001 | 8.5-10 Hz, 69.75-72.25 Hz, 72.75-105.5 Hz, 106.75-128 Hz |

**Supplementary Table T3. Comparison of EEG spectra from L6b silenced and control animals across the full frequency range recorded.** Comparisons were made between spectra 0.0-128 Hz in 0.25 Hz bins, after exclusion of bins 0.0-0.5 Hz and 49-51.5 Hz (electrical noise), using two-way ANOVAs. Results are shown for Genotype x Frequency interaction. Frontal EEG, controls n=7, L6b silenced n=9; Occipital EEG, controls n=6, L6b silenced n=9.
